# Complete Reference Genome and Pangenome Expand Biologically Relevant Information for Genome-Wide DNA Methylation Analysis Using Short-Read Sequencing and Array Data

**DOI:** 10.1101/2024.10.07.617116

**Authors:** Zheng (Joe) Dong, Joanne Whitehead, Maggie Fu, Julia L. MacIsaac, David H. Rehkopf, Luis Rosero-Bixby, Michael S. Kobor, Keegan Korthauer

**Affiliations:** Centre for Molecular Medicine and Therapeutics, BC Children’s Hospital Research Institute, University of British Columbia, 950 West 28th Avenue, Vancouver, BC, V5Z 4H4, Canada; Genome Science and Technology Graduate Program, University of British Columbia, 100-570 West 7th Avenue, Vancouver, BC, V5Z 4S6, Canada; Division of General Medical Disciplines, Department of Medicine, School of Medicine, Stanford University, 1070 Arastradero Road, Suite 300, Palo Alto, CA, 94304, USA; Centro Centroamericano de Población (CCP), Universidad de Costa Rica, San José 2060, Costa Rica; Department of Medical Genetics, University of British Columbia, 950 West 28th Avenue, Vancouver, BC, V5Z 4H4, Canada; Edwin S.H. Leong Healthy Aging Program, University of British Columbia, 117-2194 Health Sciences Mall, Vancouver, BC, V6T 1Z3, Canada; Department of Statistics, University of British Columbia, 2207 Main Mall, Vancouver, BC, V6T 1Z4, Canada

**Keywords:** T2T-CHM13, pangenome, epigenetics, bisulfite sequencing, cross-reactive probe, Illumina Infinium HumanMethylation BeadChips annotation, HumanMethylation450 BeadChip (HM450K), Illumina HumanMethylationEPIC BeadChip (EPIC), Infinium MethylationEPIC v2.0 (EPICv2), epigenome-wide association study (EWAS), cancer

## Abstract

**Background:** The new complete telomere-to-telomere human genome assembly, T2T-CHM13, and the first draft of the human pangenome reference provide unique opportunities to update the reference genome for epigenetics investigations and clinical research. However, it is largely unclear how these reference genome updates may impact DNA methylation (DNAm) analysis.

**Results:** Compared to the previous GRCh38 assembly, we found an average increase of 7.4% (range 5.4%–9.9% across samples and sequencing methods) in the number of CpGs genome-wide using T2T-CHM13 with data from four commonly used short-read sequencing DNAm profiling methods. The increase in number of CpGs facilitated discovery of 88 new differentially methylated CpGs within cancer driver genes in an epigenome-wide association study (EWAS) of colon cancer. Further, by aligning probe sequences from the commonly used and recently released Illumina DNAm arrays to T2T-CHM13 and GRCh38, we showed the enhanced utility of T2T-CHM13 for evaluation of potential probe cross-reactivity (i.e., where probes match multiple regions) and mismatch (i.e., where probes do not perfectly match the target region), resulting in the identification of new and more reproducible sets of unambiguous probes (i.e., probes uniquely mapping to the target region) (HM450K, *n* = 430,719; EPIC, *n* = 777,491; EPICv2, *n* = 859,216). In EWASs of 24 cancer types, an average of 945 additional differentially methylated CpG sites were identified in the new unambiguous probe set rather than in the GRCh38-based unambiguous probe set, with enrichments in cancer driver genes and cancer signaling pathways. Moreover, the pangenome called 4.5% more CpGs on average in short-read sequencing data than T2T-CHM13 and identified cross-population and population-specific unambiguous probes in DNAm arrays, owing to its improved representation of genetic diversity. These additional CpGs were overlapped with the promoters and gene bodies of various biologically and medically relevant genes and pangenome-based unambiguous probes can potentially facilitate the discovery of DNAm alterations in more than 200 cancer driver genes in each cancer type.

**Conclusions:** Use of T2T-CHM13 and pangenome references can benefit epigenome-wide association studies by including CpGs previously unobserved in short-read sequencing data and by improving the identification of unambiguous probes for DNAm arrays, thus expanding biologically relevant information. This study highlights the practical applications of T2T-CHM13 and pangenome for genome biology and provides a basis for expansion of epigenetics investigations.

## Background

The availability of a high-quality reference genome greatly facilitates rigorous genetic and epigenetic studies. The recently generated complete telomere-to-telomere human genome assembly, T2T-CHM13, corrects thousands of structural errors and adds nearly 200 million base pairs (bp) of sequence compared to the previous incomplete reference genome, GRCh38 ^1,2^. The previously unresolved sequences contained a large number of regions amenable to DNA methylation (DNAm), such as cytosine-phosphate-guanine (CpG) islands, which are defined as genomic regions ≥ 200 bp with GC content > 50% and observed vs. expected CpG ratio > 0.6 ^1,2^. In mammals, CpGs are the most common sites of DNA methylation, a key epigenetic mark associated with tissue development, transcriptional regulation, environmental exposures, and disease susceptibility^3,4^. The availability of the T2T-CHM13 assembly provides a timely opportunity to update the reference genome for epigenetics investigations.

A previous work suggested the major advances of T2T-CHM13 in long-read nanopore data by adding 3.6 million CpG sites (corresponding to 12.4% of total CpG sites, omitting the Y chromosome)^2^. However, only one sample sequenced by one of the short-read sequencing technologies, whole-genome bisulfite sequencing (WGBS), was examined in the previous study^2^. The potential advantages of T2T-CHM13 for DNAm analysis based on short-read sequencing DNAm profiling and array data, which are cost-effective and used extensively in current genome-wide DNAm studies, were largely undescribed.

An increasing number of studies have proven that a single linear reference genome is insufficient to represent the genetic diversity of our species, particularly due to the between-population genetic structures and within-population structural variants, and human pangenome references constructed from a diverse set of individuals are gaining popularity as a tool to capture global genomic diversity^5,6^. For instance, T2T-CHM13 v2 is derived from the CHM13 (all 22 human autosomes and chromosome X) and HG002 (chromosome Y) cell lines, both of European origin^1^. A first draft of the human pangenome reference was recently released by the Human Pangenome Reference Consortium (HPRC), containing 94 haplotypes from a diverse group of 47 individuals^7^. It added 1,115 gene duplications and 119 million base pairs of euchromatic polymorphic sequences, the majority of which were derived from structural variation, in comparison to GRCh38 ^7^. The new pangenome reference has been successfully used to identify new clinically relevant structural variants and proved to improve the calling accuracy of single-nucleotide polymorphisms (SNPs) in short-read sequencing data compared to GRCh38^7^. However, the benefit of this pangenome reference for DNAm studies remains elusive.

Due to the improvements in these updated reference genomes, we hypothesized that T2T-CHM13 and pangenome have practical advantages for DNAm analysis. We found that T2T-CHM13 has broad benefits for DNAm analysis compared to GRCh38, including increased number of CpG sites called from short-read sequencing DNAm data and reduced potential impacts of probe cross-reactivity and mismatch for DNAm array data. More importantly, these additional CpGs and unambiguous probes benefited from the use of T2T-CHM13 offer additional biological insights for epigenome-wide association studies (EWASs), even in publicly available data. We also showed that a first draft of the human pangenome reference can provide a large number of additional CpGs not present in a single linear reference genome in short-read sequencing data and can identify cross-population and population-specific unambiguous probes in DNAm arrays as a result of its improved representation of human genetic diversity. These additional CpGs and unambiguous probes could significantly advance the discovery of biologically and medically relevant DNAm alterations. Consequently, approaches to epigenetics analyses need to take these reference genome updates into account and consider their biologically relevant information expansion. This thinking underscores the importance of understanding that the epigenetic architecture of diseases using T2T-CHM13 and pangenome will have implications for future clinical approaches to disease diagnostics, prognostics, and treatment.

## Results

### T2T-CHM13 reference expanded the number of CpGs profiled by short-read DNAm sequencing

Given the increased number of CpGs in the new genome assembly, we expected that there would be an increase in CpG calls from short-read DNAm sequencing data using T2T-CHM13 compared to GRCh38, just as in long-read sequencing^2^. However, in contrast to long-read sequencing that produces long reads (usually 10–30 kb), short-read sequencing produces short reads (usually 75-300 bp)^8^. As long reads are expected to have an advantage compared to short reads in uniquely mapping to repeat regions that make up a large portion of those resolved by T2T-CHM13, we quantified how many additional CpGs can be called in short-read DNAm sequencing data using T2T-CHM13. We performed an empirical study of four commonly used short-read sequencing DNAm profiling methods: two methods utilizing bisulfite conversion, i.e., whole genome bisulfite sequencing (WGBS)^9^ and reduced-representation bisulfite sequencing (RRBS)^10^; and two alternative methods utilizing enrichment of methylated DNA, i.e., methyl-binding domain sequencing (MBD-seq)^11^ and methylated DNA immunoprecipitation sequencing (MeDIP-seq)^12^.

Across 15 cell line samples from four biosamples (i.e., human H1, H9, GM12878, and K562 cell lines) sequenced by multiple short-read sequencing methods (data sources described in Table S1), we found an average of 7.4% (range 5.4%–9.9% across samples and sequencing methods) more CpGs called genome-wide using T2T-CHM13 than GRCh38 as the reference genome (Fig. 1a, 1b, Table S1). This increase matched well with previous results of chromatin immunoprecipitation sequencing (ChIP-seq) analysis, where 3.0%–19.4% more peaks were called for seven other epigenetic marks (H3K9me3, H3K27me3, H3K27ac, H3K36me3, H3K4me1, H3K4me3, and CTCF) using T2T-CHM13 ^2^. Of CpGs called uniquely using T2T-CHM13 (i.e., additional CpGs called using T2T-CHM13 vs. GRCh38), 9.4%–54.0% were found in sequences not included in GRCh38, and others may benefit from the superiority of T2T-CHM13 in short-read mapping (i.e., one CpG in both reference genomes was only successfully called from short reads when using T2T-CHM13) (Table S2). This may be underlined by the observation that the majority of T2T-CHM13-unique CpGs (73.9%–94.6%) overlapped with segmental duplications and/or RepeatMasker-annotated repetitive regions that were corrected and expanded in the T2T-CHM13 assembly^1,13–15^ (Fig. 1c). In addition, T2T-CHM13 has been reported to be more accurate in methylation calls in repetitive regions in alignments compared to GRCh38 ^16^.

**Fig. 1.**
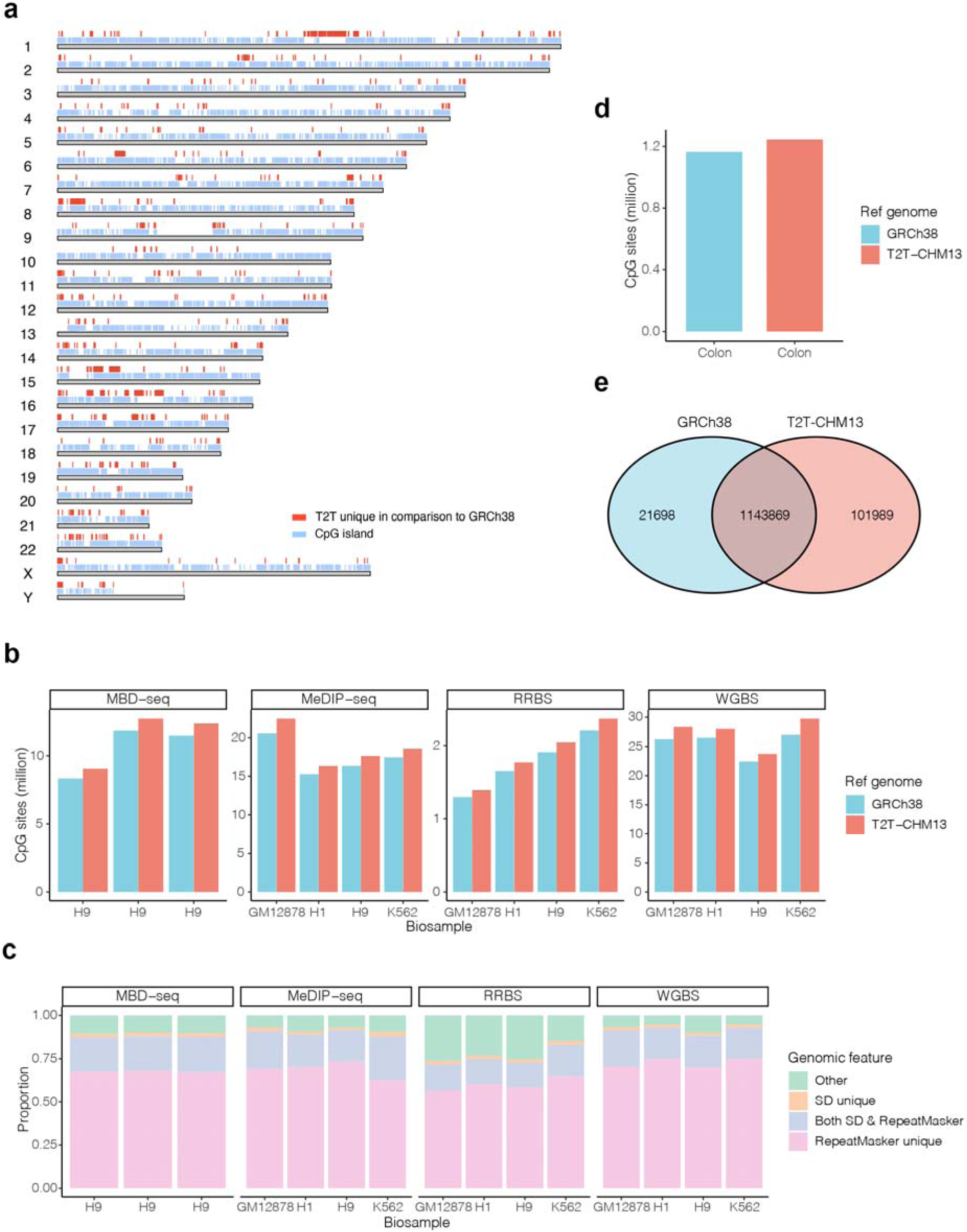
Short-read sequencing DNAm profiling using T2T-CHM13. **a** Ideogram of T2T-CHM13 v2.0 reference genome, showing T2T-unique genomic regions in comparison to GRCh38 (red) and CpG islands in T2T-CHM13 (pale blue) in a K562 WGBS sample. **b** Numbers of CpGs detected using the T2T-CHM13 (red) and GRCh38 (blue) reference genomes. The number of actual CpGs within sequencing reads called from MBD-seq and MeDIP-seq samples was determined by intersecting the genomic coordinates for the reads with all CpG coordinates in the reference genome. The three MBD-seq samples for the H9 biosample from SRR500520, SRR500521, and SRR500522 are shown from left to right. **c** The distribution of CpGs that were called uniquely using T2T-CHM13 in repetitive elements or other regions. These elements included segmental duplications (SDs) and RepeatMasker-annotated repetitive regions. **d** Numbers of CpGs detected using the T2T-CHM13 (red) and GRCh38 (blue) reference genomes in a colon cancer EWAS using RRBS data. **e** Venn diagram of CpGs identified using T2T-CHM13 and GRCh38 reference genomes in a colon cancer EWAS using RRBS data.

Moreover, 4.6%–8.7% more CpGs shared across the samples for each short-read sequencing method were called by T2T-CHM13 than with GRCh38 (Table S3). These observations suggested that most of the additional CpGs called by T2T-CHM13 represented a consistent set across multiple samples, supporting the validity of these additional CpGs in the analysis of DNAm patterns across samples. As a consequence, using T2T-CHM13 as the reference genome in short-read sequencing DNAm profiling yielded significant expansion of CpGs over GRCh38.

### EWAS detected an increased number of significantly associated CpGs using T2T-CHM13

The increases in number of CpGs using T2T-CHM13 can potentially improve analyses and facilitate the discovery of new differentially methylated CpGs in EWASs. As an example, we reanalyzed a colon cancer EWAS including RRBS samples (*n* = 20) from 10 colon tumors and 10 adjacent normal tissues, as DNAm has been recognized as a suitable biomarker for colon cancer screening^17,18^ (Table S4). Here we employed an RRBS dataset as an example due to its minimal CpG increase using T2T-CHM13 (Fig. 1b), facilitating the extension of the findings to datasets using other short-read sequencing platforms. Based on T2T-CHM13 and GRCh38, 1,245,858 and 1,165,567 CpGs were called across these samples and included in the analysis, respectively (see Methods) (Fig. 1d). Consistent with the findings outlined above, 80,291 (6.9%) more CpGs were called using T2T-CHM13 compared with GRCh38. To evaluate the potential value of these additional CpGs in the analysis of DNAm patterns in EWAS, we identified intersections (*n* = 1,143,869) between CpGs called by the two reference genomes and identified CpGs called exclusively using T2T-CHM13 (*n* = 101,989) by excluding all intersecting CpGs from the CpGs called using T2T-CHM13 (Fig. 1e).

Using thresholds of false discovery rate (FDR) < 0.05 (corresponding to permutation *P* = 3.25 × 10^−4^ and permutation *P* = 2.06 × 10^−4^ for CpGs called by T2T-CHM13 and GRCh38, respectively) and DNAm change > 0.1, 984 and 499 CpGs called by T2T-CHM13 and GRCh38 were associated with colon cancer, respectively (Table S5). The chance of identifying colon cancer-associated alterations was greater using T2T-CHM13 than GRCh38 (0.08% vs. 0.04%, respectively, Chi-squared test *P* = 1.49 × 10^−29^). Among these colon cancer-associated CpGs, 517 and 32 were uniquely identified using T2T-CHM13 and GRCh38, respectively. The 32 colon cancer-associated CpGs uniquely discovered using GRCh38 failed to be identified using T2T-CHM13 because six CpGs were not called using T2T-CHM13, and because the remaining 26 CpGs had varied association *P*-values between analyses based on the two assemblies and were therefore unable to pass the FDR threshold in T2T-CHM13-based analysis. Their varied association *P*-values were driven by different DNAm levels measured by T2T-CHM13 and GRCh38, respectively. This may be explained by the large proportion of these 26 CpGs (84.4%) in repeat sequences that were well-characterized in T2T-CHM13 ^1,13–15.^ Of the 517 colon cancer-associated CpGs uniquely discovered using T2T-CHM13, 88 were among the additional CpGs uniquely called using T2T-CHM13, while others were largely attributable to the difference in FDR estimations between T2T-CHM13-based and GRCh38-based analysis (Fig. 2a).

**Fig. 2.**
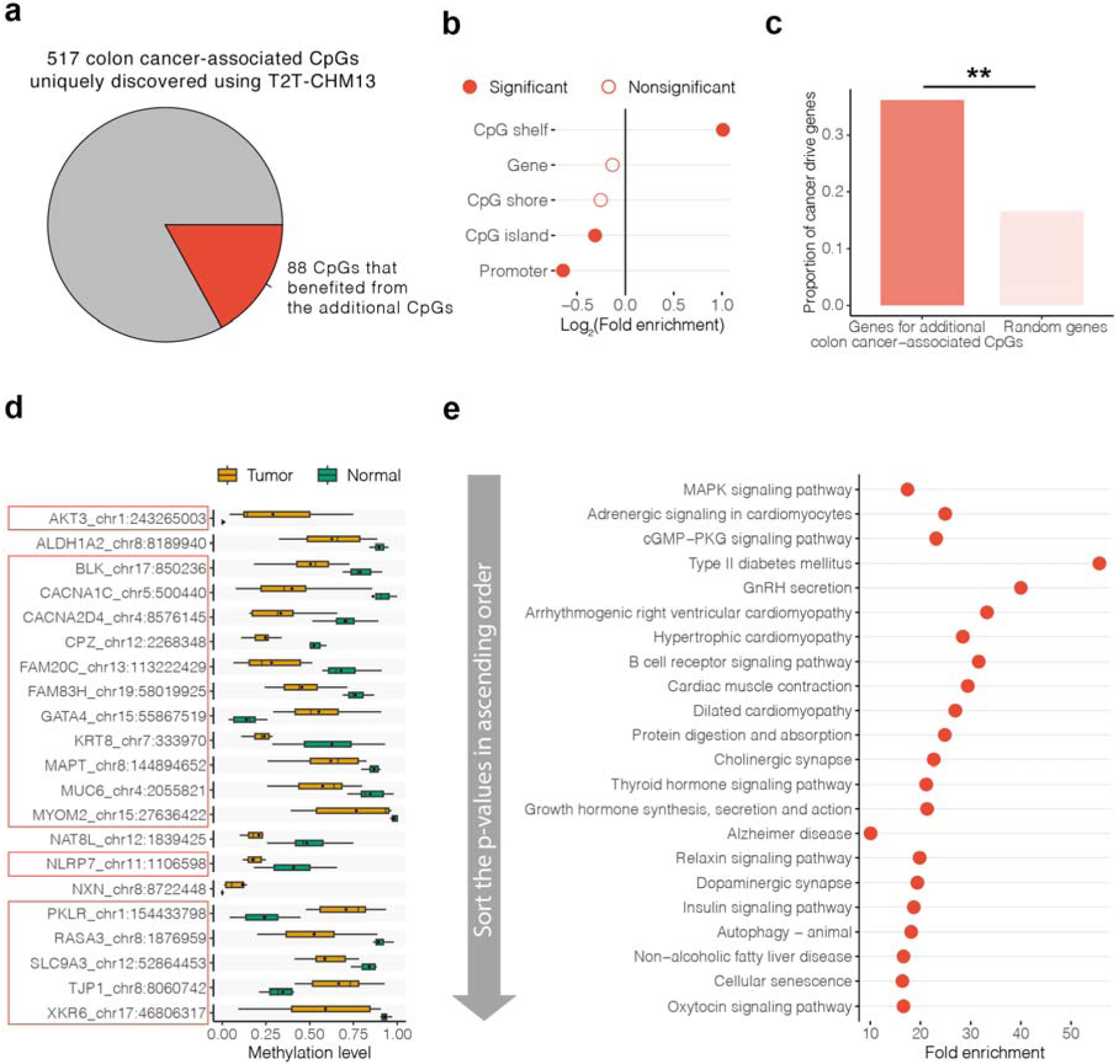
T2T-CHM13 benefits EWAS by providing additional biological insights. **a** A pie chart showing the percentage of the 88 colon cancer-associated CpGs that benefited from the additional CpGs called using T2T-CHM13 across the 517 colon cancer-associated CpGs uniquely discovered using T2T-CHM13. CpGs with FDR < 0.05 and DNAm changes > 0.1 between colon tumor (*n* = 10) and adjacent normal RRBS samples (*n* = 10) were considered as colon cancer-associated CpGs. **b** Enrichment of colon cancer-associated CpGs that benefited from the additional CpGs called using T2T-CHM13 (*n* = 88) with genomic elements. **C** Proportion of cancer driver genes in genes for the 88 colon cancer-associated CpGs that benefited from the additional CpGs called using T2T-CHM13 compared to random sets of genes. Random sampling was repeated 1000 times. ** 0.001 < *P* < 0.01. **d** Methylation patterns of additional colon cancer-associated CpGs in cancer driver genes between colon tumors (*n* = 10) and normal tissues (*n* = 10). The red bar shows the cancer driver genes, DNAm alterations of which were uniquely found by these additional colon cancer-associated CpGs. Mean values are presented as circles in each box plot. Outliers are not shown in box plots. **e** KEGG pathways for genes whose promoters and gene bodies are overlapped with the 88 colon cancer-associated CpGs. These pathways are sorted by enrichment *P*-values.

With regard to the 88 colon cancer-associated CpGs amongst the set of additional CpGs called using T2T-CHM13, we performed principal component analysis (PCA) to summarize their DNAm patterns across samples. Tumor and normal tissue samples showed distinct loadings of the top two principal components (PCs), which together accounted for 76.2% of the total variance (Fig. S1). These results suggested that T2T-CHM13 assembly can promote detection of additional CpGs associated with colon cancer by expanding the repertoire of test CpGs in EWASs utilizing short-sequencing data.

### T2T-CHM13 provided additional biological insights for the colon cancer EWAS

Identification of additional colon cancer-associated CpGs holds great promise to provide more biological insights and deepen our understanding of the role of DNAm in the pathogenesis of colon cancer. To test this, we characterized the possible functional roles of the additional 88 colon cancer-associated CpGs by overlapping them to genomic features and performing enrichment analysis for cancer driver genes and biological processes. We found that 68 of the 88 colon cancer-associated CpGs amongst the set of additional CpGs uniquely called using T2T-CHM13 were located in promoters, gene bodies, CpG islands, CpG shores (2-kb flanking regions of CpG island), and/or CpG shelves (2-kb flanking regions outward from CpG shores), DNAm patterns which play dominant roles in transcriptional regulation^19–23^ (Fig. S2). These 88 CpGs had similar distributions (i.e., no significant enrichment or depletion) in these elements compared to 896 colon cancer-associated CpGs shared with GRCh38 (all Chi-squared test FDR > 0.05), but they showed significant depletion in promoters (fold enrichment = 0.6, FDR = 0.01) and CpG islands (fold enrichment = 0.8, FDR = 7.81 × 10^−3^) and enrichment in CpG shelves (fold enrichment = 2.0, FDR = 1.4 × 10^−4^) compared to random set of all CpG sites from the RRBS data (Fig. 2b, Table S6). As CpG shelf methylation has been reported to antagonize the binding of polycomb proteins that are essential for the maintenance of the repressed state of cell-type-specific genes^23,24^, our findings were consistent with an epigenetic progenitor model of cancer, in which epigenetic alterations affecting tissue-specific differentiation are the predominant mechanism by which epigenetic changes cause cancer^22,25^. These results indicated that these additional colon cancer-associated CpGs may have regulatory functions in the development of colon cancer.

As DNAm-mediated modulation may be a fundamental mechanism influencing the regulation of cancer driver genes^26^, to obtain a snapshot of the potential roles of 58 genes whose promoters and gene bodies overlapped the 88 additional colon cancer-associated CpGs, we quantified enrichment of these genes with cancer driver genes, mutations of which affect cancer progression^27^. The cancer driver gene set was collected from Integrative OncoGenomics, which is a database of the compendium of mutational driver genes across 73 tumor types^27^ and DriverDBv4, which is a multi-omics integration database for cancer driver genes across 60 tumor types^28^. We found that 21 genes were cancer driver genes, with 2.2-fold enrichment than expected by chance (*P* = 0.002, 1000 permutation test) (Fig. 2c, Table S7). For example, *GATA4* has been demonstrated to play a substantial role in controlling proliferation and acting as a regulator of the pro-inflammatory secretome in cancer^29^. Another example was *AKT3*, which is highly expressed in mesenchymal colorectal cancer cells, and active AKT3 inhibits a cyclin-dependent kinase inhibitor, p27^KIP1^, that suppresses tumorigenesis^30^. Interestingly, DNAm alterations in 18 of these cancer driver genes (including *GATA4* and *AKT3* genes mentioned above) were uniquely found by these additional colon cancer-associated CpGs (i.e., no DNAm alterations in the promoters or gene bodies of these 18 genes were identified using GRCh38-based analysis) (Fig. 2d, Table S8). This was consistent with the improvement of T2T-CHM13 in calling genetic variants for medically relevant genes^31^ and suggested that these additional colon cancer-associated CpGs may play a crucial role in the regulation of cancer driver genes, which helps to broaden our understanding of cancer biology.

We also performed gene set enrichment analysis for the 58 genes whose promoters and gene bodies overlapped the 88 additional colon cancer-associated CpGs to reveal their related biological processes. Applying a Benjamini–Hochberg FDR threshold of 0.05, these genes showed greater enrichment in 716 biological pathways compared to all genes (Table S9). The top three ranked non-root Kyoto encyclopedia of genes and genomes (KEGG) pathways were the MAPK signaling pathway (fold enrichment = 17.4, FDR = 6.63 × 10^−3^), adrenergic signaling in cardiomyocytes (fold enrichment = 24.9, FDR = 0.01), and the cGMP-PKG signaling pathway (fold enrichment = 23.1, FDR = 0.01), which are themselves or are related to cancer signaling pathways^32–34^ (Fig. 2e). These enriched pathways, comprised of the cancer driver genes identified above, implied the biological interactions of these cancer drivers. Further compared to the 707 genes whose promoters and gene bodies overlapped the 896 colon cancer-associated CpGs shared with GRCh38, the 58 genes showed greater enrichment in 31 biological pathways, with top three ranked non-root KEGG pathways of alanine, aspartate and glutamate metabolism (fold enrichment = 179.3, FDR = 0.01), retinol metabolism (fold enrichment = 89.7, FDR = 0.01), and metabolic pathways (fold enrichment = 12.8, FDR = 0.01) (Table S10). As metabolic reprogramming is a major hallmark of cancer^35,36^, our findings on the additional colon cancer-associated CpGs may be complementary to understanding the pathogenesis of colon cancer. Taken together, T2T-CHM13 benefits in providing additional biological insights for the EWAS of colon cancer. More broadly, these results show the potential for T2T-based analyses of publicly available data resource to yield new biological insights, even in short-read sequencing data.

### T2T-CHM13 reference genome identified new sets of unambiguous probes on Illumina DNA methylation arrays

To further explore the utility of T2T-CHM13 for epigenetics research, we applied the most widely used tools for characterizing genome-wide DNAm in humans, i.e., the HM450K (*n* = 485,512), EPIC (*n* = 865,859), and the recently updated EPICv2 (*n* = 936,190) DNAm arrays, designed to measure DNAm level by the specific hybridization of each probe to the target DNAm site. Due to the sequence dependence of hybridization to these DNAm arrays, probe technical artifacts, such as cross-reactivity (also known as cross-hybridization where probes match multiple regions of a reference genome) and mismatch (i.e., where probes do not perfectly match the target region), have been shown to bias DNAm measurements and therefore affect EWAS results^37^. Cross-reactivity occurs when a probe maps to not only the target region but also to other unintended parts of the genome (Fig. 3a), potentially yielding signals that do not originate exclusively from the annotated target CpG^38–44^. A number of *in silico* investigations have identified tens of thousands of cross-reactive probes on the Illumina DNAm arrays of HM450K and EPIC^38–45^. Furthermore, probes of HM450K and EPIC arrays were designed based on the GRCh37/hg19 reference genome assembly, and a subset show mismatch when aligned to GRCh38, including the inability of some probes to map to GRCh38 and differences in the extension base of some probes between GRCh37 and GRCh38 assemblies, suggesting the impact of the reference genome update on the array probe evaluation process^43^.

**Fig. 3.**
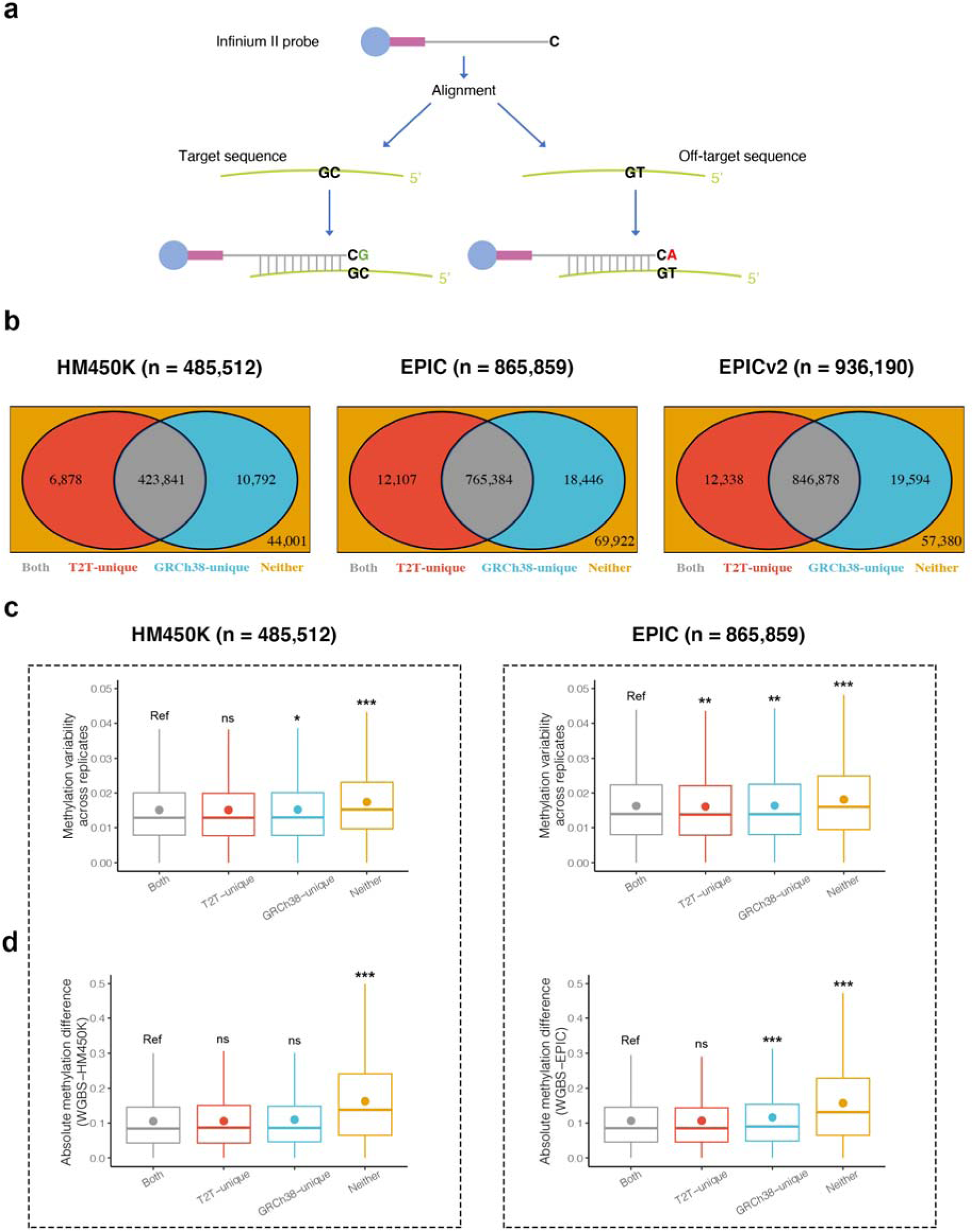
T2T-CHM13 improves the discovery of unambiguous probes (i.e., non–cross-reactive and non-mismatched probes uniquely mapping to the target region). **a** Schematic representation of the cross-reactivity of one probe. **b** Venn diagram of unambiguous probes identified using T2T-CHM13 and GRCh38 reference genomes. Other probes (“Neither”) were those with mapping problems defined by both reference genomes, including cross-reactive and/or mismatched probes. **c** DNAm variability of CpGs across technical replicates in all CpG groups for the HM450K (left, *n* = 4) and EPIC (right, *n* = 3) arrays. Mean values are represented as circles in each box plot. Not significant (ns); FDR ≥ 0.05; * 0.01 ≤ FDR < 0.05; ** 0.001 ≤ FDR < 0.01; *** FDR < 0.001. Outliers are not shown in box plots. **d** Differences in DNAm levels calculated for CpG groups between one array and one matching WGBS.

An increasing number of studies have proven that cross-reactive and mismatched probes may yield inaccurate or misleading results, and their removal is often recommended in EWASs to focus on unambiguous probes (i.e., non–cross-reactive and non-mismatched probes uniquely mapping to the target region)^38–44^. Although a number of *in silico* investigations have been performed for HM450K and EPIC arrays, and EPICv2 was designed to remove probes with these potential technical artifacts based on one of these investigations^43^, previous investigations used incomplete reference genomes (e.g., GRCh37/hg19 and GRCh38/hg38) for the identification and exclusion of such cross-reactive and mismatched probes. As T2T-CHM13 represents a gap-free reference genome and corrects structural errors in GRCh38, we performed comprehensive *in silico* analyses using this reference genome to examine whether it provides benefits in evaluating cross-reactive and mismatched probes, potentially resulting in a more reproducible set of unambiguous probes that align to specific single regions of the genome.

By *in silico* alignment using T2T-CHM13, we found 88.7% (*n* = 430,719), 89.8% (*n* = 777,491), and 91.8% (*n* = 859,216) unambiguous probes on the HM450K, EPIC and EPICv2 arrays, respectively, compared to 89.5% (*n* = 434,633), 90.5% (*n* = 783,830), and 866,472 (92.6%), respectively, using GRCh38 (Fig. 3b; The list of unambiguous probes is available for download from https://github.com/functionalepigenomics/Illumina_Infinium_HumanMethylation_BeadChips_A nnotation). Although the vast majority of probes were found to be unambiguous using both reference genomes (HM450K, *n* = 423,841; EPIC, *n* = 765,384; EPICv2, *n* = 846,878), 17,670 (3.6%), 30,553 (3.5%), and 31,932 (3.4%) probes on the HM450K, EPIC, and EPICv2 arrays, respectively, were annotated differently between reference genomes (Fig. 3b). Among these probes, most (HM450K, *n* = 10,792; EPIC, *n* = 18,446; EPICv2, *n* = 19,594) were uniquely identified as unambiguous in GRCh38, with a smaller proportion identified as unambiguous only in T2T-CHM13 (HM450K, *n* = 6878; EPIC, *n* = 12,107; EPICv2, *n* = 12,338). Therefore, the number of identified unambiguous probes was consistently smaller using T2T-CHM13 than GRCh38 for the three arrays. Similar results were observed in comparison with previous investigations using incomplete reference genomes, where previously identified unambiguous probes outnumbered those defined here using T2T-CHM13 ^39–45^ (Table S11). These findings showed that T2T-CHM13 enables a revised set of unambiguous probes.

### Unambiguous probes identified by T2T-CHM13 were more reproducible than those identified by GRCh38

To elucidate the benefits of unambiguous probe detection using T2T-CHM13 compared to GRCh38, we examined the probes with differing annotations. Here we focus on the HM450K and EPIC arrays, as their abundant datasets are available and EPIC can cover most (77.63%) of the CpGs in EPICv2. As stated in the previous section, 6878 and 12,107 probes were uniquely identified as unambiguous on the HM450K and EPIC arrays, respectively, by T2T-CHM13 (i.e., “T2T-unique” unambiguous probes) compared with GRCh38 (Table 1, Fig. 3b, Table S12). Of these probes, 16.1% (HM450K, *n* = 1110) and 15.5% (EPIC, *n* = 1878) mapped to multiple regions in GRCh38, while others were mismatched (Table 1). The discovery of these T2T-unique unambiguous probes could be a result of the correction of structural errors in GRCh38 by T2T-CHM13 ^1^. To test whether DNAm measurements were as reproducible for these T2T-unique unambiguous probes as for unambiguous probes defined by both reference genomes (i.e., “Both” unambiguous probes), we examined the variability of these probes across technical array replicates and difference in matched WGBS measurements (currently regarded as the “gold-standard” technique for quantifying DNAm), both from the lung-derived human IMR-90 cell line. This investigation was motivated by reports that probe cross-reactivity and mismatch can introduce bias, and may therefore cause poor reproducibility of probes^38–44^. This was confirmed in our analysis—probes with cross-reactivity or mismatch defined by both reference genomes (i.e., “Neither” unambiguous probes) exhibited lower reproducibility (as measured by an increase in variability, in units of standard deviation) than Both unambiguous probes across array technical replicates as well as lower absolute difference between array and matching WGBS samples (average 0.020 vs. 0.016, respectively, in technical replicates and average 0.169 vs. 0.107, respectively, between array and matching WGBS data in HM450K; and average 0.021 vs. 0.018, respectively, in technical replicates and average 0.165 vs. 0.108, respectively, between array and matching WGBS data in EPIC; all Mann–Whitney U test FDR < 0.001) (Fig. 3c, d yellow vs. gray, respectively).

**Table 1.** Category of array probes based on *in silico* alignment using T2T-CHM13 and GRCh38.

| Platform | Category | Num. in each category | Unambiguous |  | Cross-reactive |  | Mismatched |  |
| --- | --- | --- | --- | --- | --- | --- | --- | --- |
|  |  |  | T2T-CHM13 | GRCh38 | T2T-CHM13 | GRCh38 | T2T-CHM13 | GRCh38 |
| HM450K<br>(*n* = 485,512) | Both | 423,841 | 423,841 | 423,841 | 0 | 0 | 0 | 0 |
|  | T2T-unique | 6878 | 6878 | 0 | 0 | 1110 | 0 | 5768 |
|  | GRCh38-unique | 10,792 | 0 | 10,792 | 1234 | 0 | 9558 | 0 |
|  | Neither | 44,001 | 0 | 0 | 40,810 | 40,932 | 3191 | 3069 |
| EPIC<br>(*n* = 865,859) | Both | 765,384 | 765,384 | 765,384 | 0 | 0 | 0 | 0 |
|  | T2T-unique | 12,107 | 12,107 | 0 | 0 | 1878 | 0 | 10,229 |
|  | GRCh38-unique | 18,446 | 0 | 18,446 | 1861 | 0 | 16,585 | 0 |
|  | Neither | 69,922 | 0 | 0 | 66,711 | 66,871 | 3211 | 3051 |
| EPICv2<br>(*n* = 936,190) | Both | 846,878 | 846,878 | 846,878 | 0 | 0 | 0 | 0 |
|  | T2T-unique | 12,338 | 12,338 | 0 | 0 | 1113 | 0 | 11,225 |
|  | GRCh38-unique | 19,594 | 0 | 19,594 | 1768 | 0 | 17,826 | 0 |
|  | Neither | 57,380 | 0 | 0 | 54,446 | 54,536 | 2934 | 2844 |

Probes in each array were first aligned to the reference genome to check whether they had good alignments (i.e., sequence alignment with at least 90% identity, at least 40 of 50 matching bases, no gaps, and the CpG locus had to be perfectly matched); if not, they were classified as mismatched probes; if yes, they were further checked to determine whether they were unique alignments; if yes, they were classified as unambiguous (i.e., non-cross-reactive and non-mismatched) probes; otherwise, they were classified as cross-reactive probes.
For each array, probes were categorized based on their unambiguity (i.e., non-cross-reactive and non-mismatched) as determined by GRCh38 and T2T-CHM13:
Both: unambiguous probes defined by both reference genomes.
T2T-unique: unambiguous probes uniquely identified by T2T-CHM13.
GRCh38-unique: unambiguous probes uniquely identified by T2T-CHM13.
Neither: probes with mapping problems (cross-reactivity or mismatch) defined by both reference genomes.
For Neither unambiguous probes, 40,775 for HM450K, 66,664 for EPIC, and 54,409 for EPICv2 were cross-reactive probes demonstrated by both reference genomes, and 3034 for HM450K, 3004 for EPIC, and 2807 for EPICv2 were mismatched probes demonstrated by both reference genomes.

We next compared the reproducibility of T2T-unique and Both unambiguous probes among technical replicates and found that they showed comparable DNAm variability in HM450K (0.017 vs. 0.016, respectively, Mann–Whitney U test FDR = 1.00) and even slightly lower DNAm variability in EPIC (1.80 × 10^−2^ vs. 1.84 × 10^−2^, respectively, Mann–Whitney U test FDR = 3.82 × 10^−3^) (Fig. 3c red vs. gray, respectively). T2T-unique unambiguous probes exhibited similar absolute DNAm differences between array and WGBS measurements compared to Both unambiguous probes (0.107 vs. 0.107, respectively, in HM450K and 0.108 vs. 0.108, respectively, in EPIC; both Mann–Whitney U test FDR > 0.05) (Fig. 3d red vs. gray, respectively). In addition, one whole-blood sample with six replicates measured by the EPIC array and two publicly available HM450K array data sets containing 36 pairs of technical replicates in blood and cerebellum further confirmed the similar reproducibility of T2T-unique unambiguous probes to Both unambiguous probes (all Mann–Whitney U test FDR > 0.05) (Fig. S3, S4 red vs. gray, respectively). These findings showed that additional unambiguous probes specified by T2T-CHM13 were as reproducible as Both unambiguous probes, thus further supporting their “unambiguous” designation.

In contrast, thousands of unambiguous probes were also uniquely identified in GRCh38 (i.e., “GRCh38-unique” unambiguous probes) (HM450K, *n* = 10,792; EPIC, *n* = 18,446), and we next investigated their reproducibility as described above for T2T-unique unambiguous probes (Table 1, Fig. 3b, Table S13). In comparison to Both unambiguous probes in the IMR-90 cell line, these GRCh38-unique unambiguous probes tended to be slightly more variable across technical replicates (0.018 vs. 0.016, respectively, in HM450K and 0.020 vs. 0.018, respectively in EPIC; HM450K Mann–Whitney U test FDR = 0.05; EPIC Mann-Whitney U test FDR = 1.35 × 10^−3^) (Fig. 3c blue vs. gray, respectively), and had slightly larger absolute DNAm differences between array and matching WGBS data, although they were statistically significant only for the EPIC array (0.110 vs. 0.107, respectively, in HM450K and 0.120 vs. 0.108, respectively, in EPIC; HM450K Mann–Whitney U test FDR = 0.50; EPIC Mann–Whitney U test FDR = 5.76 × 10^−5^) (Fig. 3d blue vs. gray, respectively). While the difference across technical replicates was statistically significant, it should be noted that the effect size was extremely small. Therefore, we further confirmed these findings using the whole-blood EPIC and blood and cerebellum HM450K array data sets mentioned above. We found similar results for GRCh38-unique unambiguous probes, which had greater variability than Both unambiguous probes (all Mann– Whitney U test FDR < 0.05) (Fig. S3, S4 blue vs. gray, respectively). Therefore, some of the GRCh38-unique unambiguous probes may exhibit cross-reactivity or mismatch, consistent with our results of analyses using T2T-CHM13 showing that they were cross-reactive or mismatched probes (Table 1).

Finally, to further evaluate the potential cross-reactivity of GRCh38-unique unambiguous probes, we examined their enrichment in repetitive regions. Our analyses showed that 71.89% (*n* = 7758) and 76.15% (*n* = 14,046) of GRCh38-unique unambiguous probes on the HM450K and EPIC arrays were found in repeat regions, such as segmental duplications and satellites (Table S14), which were slightly higher than the proportions for the T2T-unique unambiguous probes (69.61% on HM450K and 75.22% on EPIC). Compared with T2T-unique unambiguous probes in each repeat region category (see Methods), GRCh38-unique unambiguous probes were moderately enriched in segmental duplications, suggesting their potential cross-reactivity (HM450K fold = 2.37, Chi-squared test *P* < 2.20 × 10^−16^; EPIC fold = 2.28, Chi-squared test *P* < 2.20 × 10^−16^) (Table S14). This shows the capability of T2T-CHM13 to identify possible cross-reactive probes in complex segmental duplication regions, consistent with previous findings that T2T-CHM13 is nine times more predictive of human copy number than GRCh38 in nonsyntenic segmental duplications^13^.

Taken together, these findings suggested that T2T-CHM13 showed beneficial effects, not only in reducing probes with potentially lower replicability defined as unambiguous using the previous incomplete reference genome but also in enhancing detection of unambiguous probes with potentially higher replicability.

### Unambiguous CpGs uniquely called by T2T-CHM13 were associated with a wide array of cancers

To add further context to the DNAm quantified at T2T-unique unambiguous sites, we examined their biological significance. Using publicly available DNAm array datasets that cover 24 cancer types (n = 8407, data sources described in Table S15), we investigated DNAm differences between tumors and normal tissues (i.e., adjacent normal tissues or healthy individual tissues) in each cancer type. We employed HM450K datasets as examples because the majority of CpG sites from HM450K are present in EPIC and EPICv2 arrays, allowing for the extension of findings to datasets using these upgraded DNAm arrays. We performed these EWASs using unambiguous probes identified by GRCh38 (*n* = 434,633) and T2T-CHM13 (*n* = 430,719) separately and detected two sets of differentially methylated CpGs (DMCs) for each of the 24 cancers using the Mann–Whitney U test with a threshold of FDR < 0.05 and DNAm change > 0.1 (see Methods).

Comparing the two sets in each cancer, an average of 945 DMCs (range from 210 to 1583) were exclusively found using T2T-CHM13, covering 13.74% (range from 3.05% to 23.02%) of unambiguous CpGs uniquely called by T2T-CHM13 (Fig. 4a). The first two PCs of these DMCs from each cancer type explained an average of 38.0% of the total variance across cancer types (Fig. S5). In total, 4263 DMCs were exclusively found using T2T-CHM13 in at least one cancer type and covered more than half (61.98%) of T2T-unique unambiguous sites, suggesting the benefit of taking account of unambiguous CpGs uniquely called by T2T-CHM13 in cancer EWASs (Fig. 4b).

**Fig. 4.**
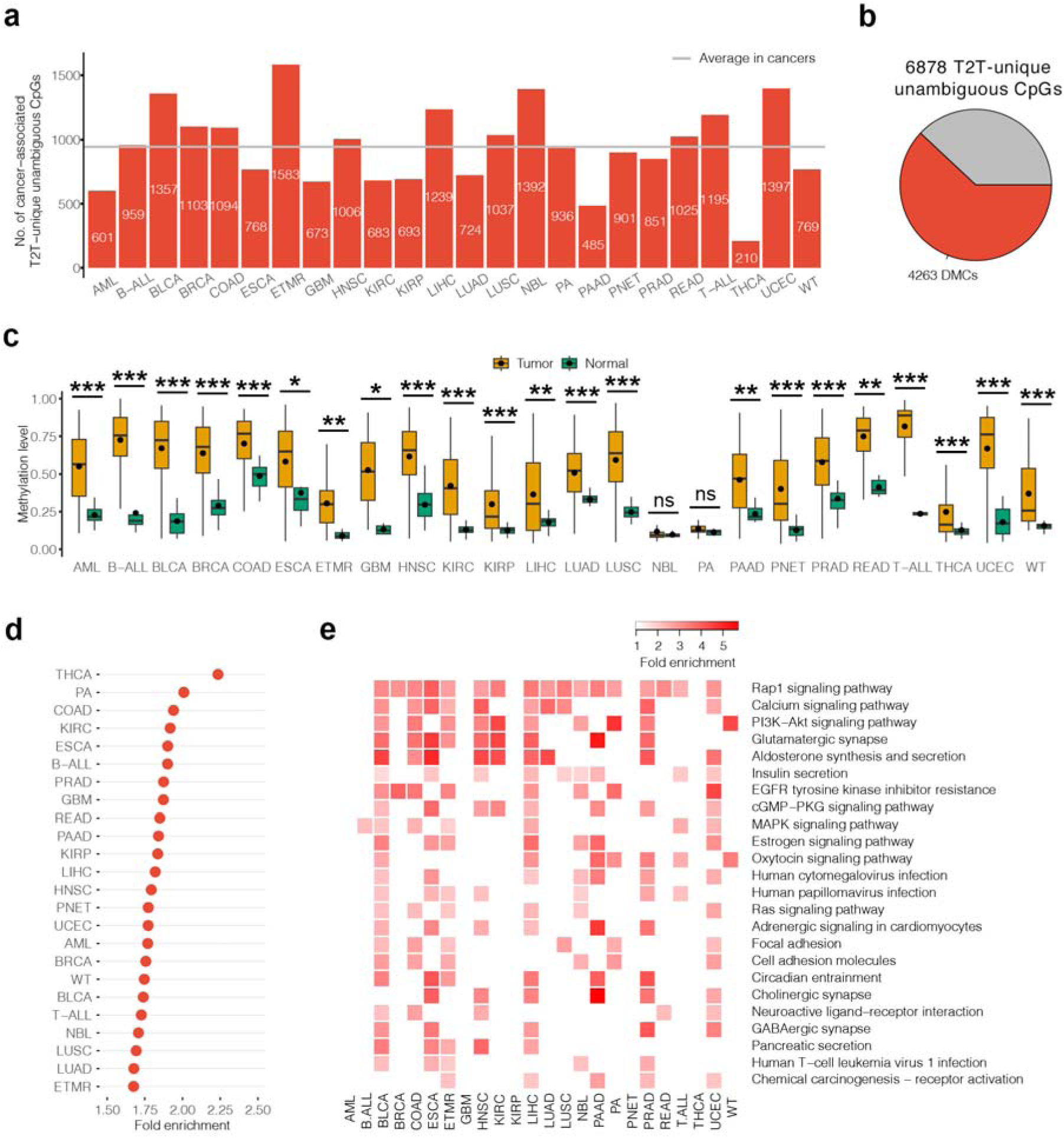
Biological significance of T2T-unique unambiguous CpGs. **a** Numbers of T2T-unique unambiguous CpGs that are differentially methylated in cancers. Differentially methylated CpGs (DMCs) are considered CpGs that are significantly associated with each cancer with a threshold of FDR < 0.05 and DNAm change > 0.1. Average number of cancer-associated T2T-unique unambiguous CpGs in 24 cancer types is shown as the gray solid line. AML, Acute myeloid leukemia. B-ALL, B-cell acute lymphoblastic leukemia. BLCA, Bladder urothelial carcinoma. BRCA, Breast invasive carcinoma. COAD, Colon adenocarcinoma. ESCA, Esophageal carcinoma. ETMR, Embryonal tumors with multilayered rosettes. GBM, Glioblastoma. HNSC, Head and neck squamous cell carcinoma. KIRC, Kidney renal clear cell carcinoma. KIRP, Kidney renal papillary cell carcinoma. LIHC, Liver hepatocellular carcinoma. LUAD, Lung adenocarcinoma. LUSC, Lung squamous cell carcinoma. NBL, Neuroblastoma. PA, Pilocytic astrocytoma. PAAD, Pancreatic adenocarcinoma. PNET, Primitive neuroectodermal tumor. PRAD, Prostate adenocarcinoma. READ, Rectum adenocarcinoma. T-ALL, T-cell acute lymphoblastic leukemia. THCA, Thyroid carcinoma. UCEC, Uterine corpus endometrial carcinoma. WT, Wilms tumor. **b** A pie chart showing the percentage of DMCs across the T2T-unique unambiguous CpGs. **c** Methylation patterns of cg27131891 between tumors and normal tissues in each cancer. Mean values are presented as circles in each box plot. Outliers are not shown in box plots. Statistical significance is denoted separately for the comparisons of methylation levels in tumors with those in normal tissues by asterisks. * FDR < 0.05; ** FDR < 0.01; *** FDR < 0.001; ^ns^ nonsignificant. **d** Significant enrichment of genes whose promoters and gene bodies are overlapped with T2T-unique DMCs in cancer driver genes. FDR < 0.05 is considered significant. **e** Shared KEGG pathways for genes whose promoters and gene bodies are overlapped with T2T-unique DMCs in 24 cancer types. Only KEGG pathways that are found in at least five cancer types are shown.

Interestingly, 81.70% of DMCs found using T2T-unique unambiguous probes were shared with at least two cancer types, highlighting the possible common mechanisms of DNAm in the tumorigenesis of multiple cancers. Particularly, cg27131891 was found to be hypermethylated in tumors across 22 cancer types (Fig. 4c). This hypermethylated CpG is in the gene body of *AL021918.1*, DNAm alterations in which have been shown to be associated with poor overall survival in lung adenocarcinoma (LUAD) patients^46^. This discovery was followed by cg01525538, cg10954469, and cg13307880, which displayed hypermethylation in tumors across 21 cancer types and located within the gene body regions of previously reported cancer-related genes, *DUSP5P1*, *LINC01143*, and the *PCDH* gene cluster^47–49^.

### Genes whose promoters and gene bodies overlapped differentially methylated unambiguous CpGs uniquely called by T2T-CHM13 were enriched in cancer driver genes and cancer signaling pathways

To characterize the possible functional roles of DMCs found using T2T-unique unambiguous probes in each cancer, we overlapped them to genomic features, particularly promoters and CpG islands, and performed enrichment analysis for cancer driver genes and biological processes. We observed that the majority of these DMCs (mean: 91.84%, range from 87.62% to 96.66%) were located in promoters, gene bodies, CpG islands, CpG shores, and/or CpG shelves (Table S16). In comparison to randomly selected CpG sites, they showed cancer-specific depletion in promoters (fold enrichment = 0.7–0.9, all Chi-squared test FDR < 0.05) and CpG islands (fold enrichment = 0.4–0.9, all FDR < 0.05) in the majority of cancer types examined (15/24 and 17/24) and enrichment in CpG shores in kidney renal papillary cell carcinoma, liver hepatocellular carcinoma, and medulloblastoma (all fold enrichment = 1.2, all FDR < 0.05) (Table S16). In contrast to the DMCs shared with GRCh38, these T2T-unique DMCs demonstrated notable enrichment solely in CpG islands within neuroblastoma, juvenile pilocytic astrocytoma tumor, and Wilms’ tumor (fold enrichment = 1.2–1.4, all Chi-squared test FDR < 0.05); other comparisons yielded no significant findings (Table S16). These findings were generally consistent with our results observed in colon cancer RRBS data using T2T-CHM13.

Given that DNAm in promoters and gene bodies has been widely reported to be correlated with gene expression^19,20^, we overlapped the DMCs uniquely found using T2T-CHM13 to promoters and gene bodies and then investigated enrichments of these corresponding genes to cancer driver genes to investigate these DMCs’ potential biological relevance. The cancer driver gene set was described above as collected from Integrative OncoGenomics^27^ and DriverDBv4 ^28^. We found that an average of 299 of the genes with promotor- or gene body-overlapped T2T-unique DMCs (range from 88 to 455) were cancer driver genes in each cancer type, with an average 1.83-fold enrichment (range from 1.68 to 2.24) compared to that expected by chance (all FDR < 0.001, 1000 permutation test) (Fig. 4d, Table S17). Notably, as compared to genes with promotor- or gene body-overlapped shared DMCs with GRCh38, these genes also showed a significant enrichment in cancer driver genes in each cancer (average fold enrichment = 1.28, range from 1.12 to 1.43, all FDR < 0.05, 1000 permutation test) (Table S17). Although the majority of these cancer driver genes can also be identified to present DNAm alterations by using GRCh38-based analysis, an average of 21 genes (range from 16 to 29) were uniquely identified by T2T-unique unambiguous sites in each cancer, with the most common gene being *CBS* (22/24 of test cancer types). *CBS* has been previously characterized as having oncogenic roles in promoting tumor growth and progression, initiating tumor formation in colon, ovarian, and breast cancer, and as a potential target for cancer therapy^50^. These were consistent with the observed benefit of T2T-CHM13 in the discovery of cancer driver genes’ methylation alterations in colon cancer RRBS data and highlighted that DNAm alterations in cancer driver genes from unambiguous CpGs uniquely called by T2T-CHM13 may improve our understanding of cancer biology and hold possible significant clinical translational value.

We also performed enrichment analysis for the genes with promotor- or gene body-overlapped T2T-unique DMCs to reveal their related biological processes. Applying a Benjamini–Hochberg FDR threshold of 0.05, these genes showed significant enrichment in an average of 510 biological pathways (range from 205 to 823) compared to all genes that overlapped the entire set of HM450K CpGs (Table S18). The top three common non-root KEGG pathways across these cancers were the Rap1 signaling pathway, the calcium signaling pathway, and the PI3K-Akt signaling pathway, which are well-known cancer-related signaling pathways^51–53^ (Fig. 4e). Compared to genes with promotor- or gene body-overlapped shared DMCs with GRCh38, these genes showed greater enrichment in an average of 134 biological pathways (rang from 38 to 346), except for THCA. The top three common non-root KEGG pathways were Rap1 signaling, morphine addiction, and MAPK signaling (Table S18). These cancer-related pathway observations benefited from T2T-unique unambiguous sites, which may help us better understand epigenetic roles in cancer biology.

### Pangenome reference expanded the number of CpGs profiled by short-read DNAm sequencing

The pangenome reference based on 94 haplotypes of 47 genetically diverse individuals has the promise to improve DNAm studies by accounting for genetic diversity that is missing from the single linear reference genome (Fig. 5a). Here we tested whether mapping using this pangenome could improve CpG calls from short read DNAm sequencing data. We aligned read sequences of the MeDIP-seq cell line samples tested above to the pangenome graph (Minigraph-Cactus [MC graph) and called CpG sites. We found the number of CpGs called was on average 3.9% (0.9%– 7.3%) higher per sample than those using T2T-CHM13 reference genome graph (based on T2T-CMH13 v2.0 reference sequence and a variant call format [VCF] file of the 1000 genomes project recalled on T2T-CHM13 v2.0) separately (Fig. 5b, Table S19). The H9 cell line showed the smallest increase (0.9%) in CpG calls when using the pangenome. Similar results were observed in three MBD-seq H9 samples, with an average increase of 0.8% (Table S19). We can only speculate on the biological reasons that could explain the slight increment, such as the relatively high genomic similarity between the H9 and CHM13 cell lines. As these samples were all from individuals with European ancestry, to further validate our findings in other human populations, we also investigated a publicly available MeDIP-seq dataset with 13 chronic lymphocytic leukemia patients (CCL) and normal B cell samples of East Asian ancestry^54^ (Table S19). We observed a similar trend with an average increase of 4.7% (3.1%–6.1%) when using the pangenome (Fig. 5b). This trend was also consistent with the prior findings determined in other epigenetic marks, such as H3K4me1, H3K27ac, and chromatin accessibility^7^.

**Fig. 5.**
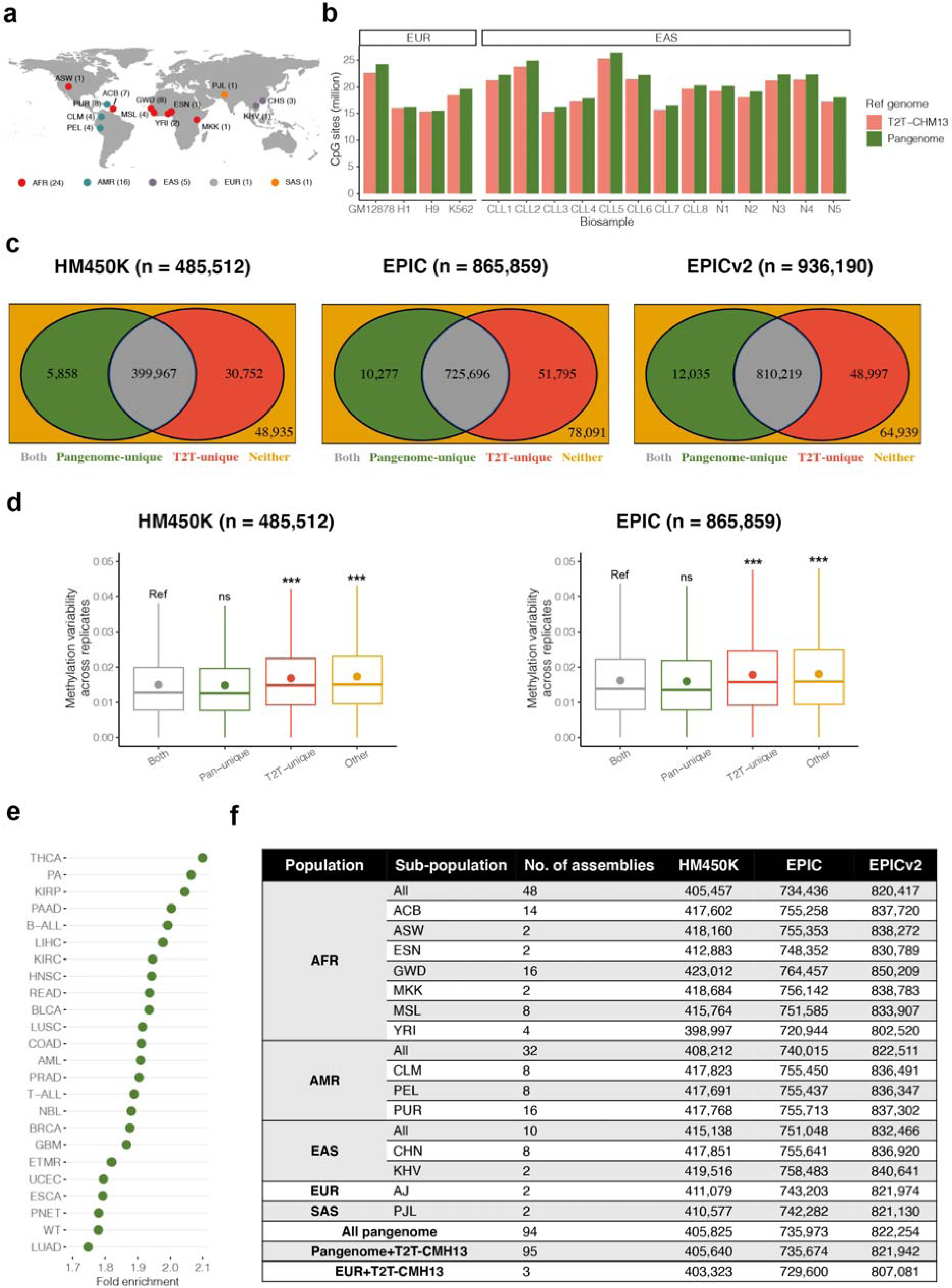
Benefits of the human pangenome in DNAm investigations. **a** World map showing the distribution of the sample populations and subpopulations in the human pangenome reference. Human populations: AFR, African; AMR, American; EAS, East Asian; EUR, European; SAS, South Asian. AFR subpopulations: ACB, African Caribbean in Barbados; ASW, African Ancestry in Southwest US; ESN, Esan in Nigeria; GWD, Gambian in Western Division; MKK, Maasai in Kinyawa, Kenya; MSL, Mende in Sierra Leone; YRI, Yoruba in Ibadan. AMR subpopulations: CLM, Colombian in Medellin, Colombia; PEL, Peruvian in Lima, Peru; PUR, Puerto Rican in Puerto Rico. EAS subpopulations: CHS, Chinese; KHV, Kinh in Ho Chi Minh City, Vietnam. EUR subpopulations: AJ, Ashkenazi Jewish. SAS subpopulations: PJL, Punjabi in Lahore, Pakistan. **b** Numbers of CpGs detected using the pangenome (green) and T2T-CHM13 (red) reference genomes. The number of actual CpGs within sequencing reads called from MeDIP-seq samples was determined by grepping “CG” from the consensus FASTA file. **c** Venn diagram of unambiguous probes identified using the pangenome and T2T-CHM13 reference genomes. Other probes were those with mapping problems defined by both reference genomes, including cross-reactive and/or mismatched probes. **d** DNAm variability of CpGs across technical replicates in all CpG groups for the HM450K (left, *n* = 4) and EPIC (right, *n* = 3) arrays. Mean values are represented as circles in each box plot. Not significant (ns); FDR ≥ 0.05; * 0.01 ≤ FDR < 0.05; ** 0.001 ≤ FDR < 0.01; *** FDR < 0.001. Outliers are not shown in box plots. **e** Significant enrichment of genes whose promoters and gene bodies are overlapped with DMCs that are exclusively found using the pangenome in cancer driver genes. FDR < 0.05 is considered significant. **f** Number of cross-population and population-specific unambiguous probes in DNAm arrays identified using the human pangenome reference.

More importantly, we found a subset of additional CpGs called by the pangenome in these samples that may illustrate DNAm patterns in medically relevant genes. For example, additional CpGs in one of these chronic lymphocytic leukemia patients (CCL3) were overlapped with promoters and gene bodies of various genes related to disease pathogenesis, such as *ABCG2*, *NRG1*, *NXN*, *PDE4D*, and *TP73*. These genes have been reported to be structurally variable in previous investigations^55^. This could be attributed to the ability of the human pangenome to represent medically relevant complex structural variants^7^. These results suggested that the pangenome can add more CpGs in EWASs utilizing short-sequencing data, thus potentially advancing the discovery of DNAm alterations of functional relevance.

### Pangenome reference aided in the identification of cross-population unambiguous probes from DNAm arrays

To create an epigenetics community resource to aid the preprocessing of array data and the downstream analyses of DNAm from individuals of different ancestries, we separately aligned probe sequences of the three DNAm arrays to each assembly of the human pangenome and identified an unambiguous probe set (see Methods). By integrating all unambiguous probe sets across 94 assemblies within the human pangenome, we identified a set of unambiguous probes suitable for DNAm testing in individuals from a diverse set of ancestries. There were 405,825, 735,973, 822,254 cross-population unambiguous probes identified in HM450K, EPIC, and EPICv2, respectively (The list of unambiguous probes is available for download from https://github.com/functionalepigenomics/Illumina_Infinium_HumanMethylation_BeadChips_Annotation).

Compared to linear reference genomes, the majority of these unambiguous probes were shared with T2T-based unambiguous probe sets (HM450K, *n* = 399,967; EPIC, *n* = 725,696; EPICv2, *n* = 810,219). The unambiguous probes identified uniquely by the pangenome (HM450K, *n* = 5858; EPIC, *n* = 10,277; EPICv2, *n* = 12,035) may benefit from accounting for genetic diversity (within-population and between-population) that is missing from T2T-CHM13 (Fig. 5c). To assess the reproducibility of these pangenome-unique unambiguous probes, we compared variability of technical array replicate and differences in matched cross-platform DNAm measurements to that of the unambiguous probes uniquely identified by T2T-CHM13. The unambiguous probes uniquely identified by the pangenome were consistently more likely to be reproducible using IMR-90 technical array replicates and WGBS measurements compared with those uniquely identified by T2T-CHM13 (0.016 vs. 0.020, respectively, in technical replicates and 0.107 vs. 0.121, respectively, between array and matching WGBS data in HM450K; and 0.018 vs. 0.021, respectively, in technical replicates and 0.109 vs. 0.123, respectively, between array and matching WGBS data in EPIC; all Mann–Whitney U test FDR < 0.001) (Fig. 5d). This result emphasizes the impact of accounting for genetic diversity when determining unambiguous probes. It is noteworthy to mention that while GRCh38 serves as a reference for genetic diversity, there were more (HM450K, fold = 6.11; EPIC, fold = 5.06; EPICv2, fold = 4.65) unambiguous probes uniquely identified by the pangenome when comparing them to GRCh38 (HM450K, *n* = 35,800; EPIC, *n* = 52,014; EPICv2, *n* = 55,976) than when comparing them to T2T-CHM13. This might suggest the significant benefits of using pangenomes for curating and evaluating ambiguous probe sets, not only by considering genetic diversity but also by ensuring a high level of reference genome completeness.

Through reanalyzing the cancer EWASs above using the unambiguous probes identified by the pangenome, we sought to elucidate the possible functional relevance of these unambiguous probes identified uniquely by the pangenome. On average, 840 DMCs (ranging from 227 to 1,314) were uniquely identified using the pangenome. These DMCs overlapped with the promoters and gene bodies of 284 cancer driver genes on average (ranging from 87 to 411), showing an average 1.91-fold enrichment (range from 1.75 to 2.10) than expected by chance (all FDR < 0.001, 1000 permutation test) (Fig. 5e, Table S20). Even compared to genes whose promoters and gene bodies overlapped the DMCs shared with T2T-CHM13, these genes also showed a significant enrichment in cancer driver genes in each cancer (average fold enrichment = 1.35, range from 1.22 to 1.48, all FDR < 0.05, 1000 permutation test) (Table S20). Furthermore, an average of 16 cancer driver genes (range from 8 to 22) were uniquely identified using the unambiguous probes identified by the pangenome (i.e., cancer-associated DNAm alterations of these genes cannot be discovered by using the T2T-based unambiguous probe set) (Table S20). Of them, the most common cancer driver gene across cancer types was *B3GNTL1* (16/24 test cancer types), DNAm alteration of which has been recently discovered through bisulfite sequencing as a component of a noninvasive biomarker for early detection of cancer^56^. These suggested the possible biological relevance of these pangenome-unique unambiguous probes in understanding cancer biology.

### Pangenome reference aided in the identification of population-specific unambiguous probes from DNAm arrays

As a large number of epigenetics studies were performed on one specific human population, ancestry-specific unambiguous probe sets were further identified at population and subpopulation levels (see Methods). At the population level, varied numbers (HM450K, *n* = 405,457–415,138; EPIC, *n* = 734,436–751,048; EPICv2, *n* = 820,417–832,466) of unambiguous probe set specific to each of five major human populations (African [AFR], American [AMR], East Asian [EAS], European [EUR], and South Asian [SAS]) were identified (Fig. 5f; The list of unambiguous probes is available for download from https://github.com/functionalepigenomics/Illumina_Infinium_HumanMethylation_BeadChips_Annotation). Despite small differences in the number of unambiguous probes across human populations, we found a subset of these population-specific unambiguous probes may provide DNAm patterns in medically relevant genes. For example, EAS-specific unambiguous probes (*n* = 484) from the HM450K array were overlapped with promoters and gene bodies of 140 cancer driver genes, such as *CTNNB1*, *DSP*, *FRK*, *HLA-A*, *NOX4*, and *TRIO* (Table S21). Consistently, some of them have been demonstrated to be associated with genetic divergences between human populations^57^.

Furthermore, the genetic structures of human subpopulations, particularly AFR subpopulations, have been widely studied^58,59^. To optimize the use of unambiguous probes in studying specific human subpopulations, varied numbers of unambiguous probe sets specific to each of the 12 subpopulations (AFR subpopulations: African Caribbean in Barbados [ACB], African Ancestry in Southwest US [ASW], Esan in Nigeria [ESN], Gambian in Western Division [GWD], Maasai in Kinyawa, Kenya [MKK], Mende in Sierra Leone [MSL], and Yoruba in Ibadan [YRI]; AMR subpopulations: Colombian in Medellin, Colombia [CLM], Peruvian in Lima, Peru [PEL], and Puerto Rican in Puerto Rico [PUR]; EAS subpopulations: Chinese [CHS] and Kinh in Ho Chi Minh City, Vietnam [KHV]) were identified (HM450K, *n* = 398,997–423,012; EPIC, *n* = 720,944–764,457; EPICv2, *n* = 802,520–850,209) (Fig. 5f; The list of unambiguous probes is available for download from https://github.com/functionalepigenomics/Illumina_Infinium_HumanMethylation_BeadChips_Annotation).

To be noted, because of the limited number (n < 5) of assemblies for populations of EUR and SAS and subpopulations of ESN, MKK, ASW, YRI, and KHV involved in the draft of the pangenome, caution should be used when using unambiguous probe sets identified for these populations and subpopulations. This limit will be greatly addressed as the human pangenome improves in the near future to capture global genomic diversity across the 700 assemblies of 350 individuals^6^.

Taken together, the pangenome reference can improve the identification of cross-population and population-specific unambiguous probes in DNAm arrays by taking advantage of genetic diversity, and thus benefiting EWASs and expanding biologically relevant information.

### Create annotation files for DNAm arrays based on T2T-CHM13 and pangenome findings

EWASs are frequently used to detect genome-wide epigenetic variants (predominantly DNAm) that are significantly associated with diseases and phenotypes of interest. However, an important concern in interpreting EWAS analyses is the potential technical bias that may arise because widely used DNAm arrays were developed based on linear incomplete reference genomes^37–44^. Assessment of technical artifacts (such as cross-reactivity and mismatch) of array probes using T2T-CHM13 and pangenome provided an opportunity to develop a community resource for the preprocessing of DNAm array data. Therefore, the unambiguous probe sets on the HM450K, EPIC, and EPICv2 arrays assessed by T2T-CHM13, pangenome, and their combination (95 assemblies including T2T-CHM13 and 94 assemblies from the pangenome; HM450K, *n* = 405,640; EPIC, *n* = 735,674; EPICv2, *n* = 821,942), separately, are available for download from https://github.com/functionalepigenomics/Illumina_Infinium_HumanMethylation_BeadChips_Annotation (Fig. 5f). Cross-population and population-specific unambiguous probes identified using the pangenome reference are also available. More importantly, we provide T2T-CHM13-based annotation information regarding promoters, gene bodies, CpG islands, CpG shores, and CpG shelves for each unambiguous probe, which may be leveraged to infer the potential biological functions of DNAm alterations in EWASs.

## Discussion

The availability of updated reference genomes is expected to significantly affect research in human genetics and epigenetics, as T2T-CHM13 has resolved complex regions^1,13–15^ and the pangenome can represent genetic diversity^5,6^. Here, we concentrated on the current most commonly used techniques for genome-wide DNAm analysis and showed that use of T2T-CHM13 and pangenome enabled the mapping of more CpG sites in short-read sequencing data and by reduced the potential ambiguities caused by probe cross-reactivity and mismatch on DNAm arrays. More importantly, we found that the increase in number of CpGs in short-read sequencing data and the identification of new unambiguous probes in DNAm arrays with use of updated reference genomes facilitated discovery of new biologically and medically relevant DNAm alterations in EWASs, with examples of cancer EWASs. These improvements may be useful in future studies and help to refine human epigenetics and epigenomics analyses.

Our study suggests that using these new reference genomes in analyses of short-read sequencing data will be valuable for the broader epigenetics community, although long-read nanopore data with the T2T-CHM13 reference genome compared to GRCh38 represents a greater benefit (12.4% more CpGs, omitting the Y chromosome) than the increase in short-read sequencing data (7.4% more CpGs on average)^2^. Nevertheless, while long-read sequencing technologies are constantly improving, short-read sequencing is currently more cost-effective and has higher per-read accuracy^8,60^. Furthermore, numerous international epigenetics projects, such as BLUEPRINT^61^, the NIH Roadmap Epigenomics Program^62^, and the Encyclopedia of DNA Elements (ENCODE)^63^, mainly used short-read sequencing methods. Therefore, this study is relevant for understanding the advantages of using T2T-CHM13 and pangenome in analysis of DNA methylomes.

Given these advantages, T2T-CHM13 and pangenome are suggested as reference genomes for calling CpGs in short-read sequencing data. It may also be worth revisiting previously published analyses using the new T2T-CHM13 and the pangenome reference genomes in certain cases. For example, given our observation that the majority of additional CpGs were annotated as repetitive regions, the additional CpGs called by T2T-CHM13 from published data sets may provide valuable additional biological information for investigations of the roles of DNAm within repetitive elements in phenotypes of interest, such as cancer and aging^64^. The pangenome can further improve CpG calling and obtain more CpGs than T2T-CHM13. It should be pointed out that the available bioinformatic tools differ between T2T-CHM13 and pangenome. T2T-CHM13 is a linear reference genome, similar to the widely used references GRCh37 and GRCh38, so conventional bioinformatic tools (e.g. Bismark^65^ and BSMAP^66^) for GRCh37 and GRCh38 can be directly applied to epigenetic analyses using T2T-CHM13. The pangenome reference is a collection of diverse genomes, so new bioinformatic tools are required and are under ongoing development. Therefore, although we found that the pangenome outperforms T2T-CHM13 in calling CpGs using currently available tools, the choice of reference genomes in epigenetics investigations should take future evaluations of newly developed pangenome mapping tools into consideration.

T2T-CHM13 and pangenome can be used to update the annotations of cross-reactive and mismatched probes in DNAm arrays, which are the most commonly used tools for characterizing genome-wide DNAm in human populations. Using T2T-CHM13, we identified a list of unambiguous probes that our analyses suggest is more reproducible than previously defined unambiguous probe lists, with improved annotation for around 3.5% of probes on the arrays. Further, cross-population and population-specific unambiguous probes in DNAm arrays were identified using the pangenome. Notably, methylation differences in some probes that were uniquely called by T2T-CHM13 or the pangenome were associated with a wide array of cancers through their possible regulatory roles in cancer driver genes and genes involved in cancer signaling pathways. As annotations in previous studies were based on incomplete reference genomes, the probes newly annotated by T2T-CHM13, the pangenome, and their combination (95 assemblies including T2T-CHM13 and 94 assemblies from the pangenome), separately, may be beneficial in preprocessing array data. This could aid in the genome-wide identification of DNAm alterations associated with diseases or variables of interest and catalyze community efforts to explore how our epigenomes influence biology and disease across human populations. Here, we have provided lists of all the newly annotated unambiguous probes in an accessible fashion as a resource for the epigenetics community.

Given the improvement in evaluation of cross-reactivity, T2T-CHM13 and the pangenome may also be effective for the design and evaluation of primers for PCR-based locus-specific DNAm analysis techniques. Similar to DNAm array probes, the primers for locus-specific techniques, such as bisulfite pyrosequencing and methylation-specific PCR, should be specific to the target DNA sequence to avoid amplification of unintended regions of the genome^67^. These PCR-based locus-specific strategies for DNAm analysis are widely used for assessment of the DNAm within specific genes or regions of interest and for validation of genome-wide DNAm analysis^68^. Therefore, the potential benefits of T2T-CHM13 and pangenome for primer design and evaluation would further expand its applicability in the field of DNAm research.

This study had the limitation that the unambiguity (i.e., non–cross-reactivity and non-mismatch) of array probes in DNAm arrays was not confirmed by experiments. As in previous studies, they were defined based on investigations against the reference genomes, and their reproducibility was characterized using technical array replicates and WGBS data^38–44^. Although the increase in CpGs called by T2T-CHM13 and the pangenome from short-read sequencing data suggested the potential for improvement of DNAm analysis, the current lack of high-quality annotation for these updated reference genomes, such as transcription factor-binding sites and chromatin states across tissues, restricts elucidation of their underlying functional significance. However, this limitation will likely be alleviated by the accumulation of data from additional studies regarding T2T-CHM13 and the pangenome, which will facilitate high-quality annotation of the human epigenome.

## Conclusions

The T2T-CHM13 and pangenome references can potentially improve EWASs by adding more CpGs in short-read sequencing data and improving the identification of unambiguous probes for DNAm arrays, thus expanding biologically relevant information. This study advanced our understanding of the practical application of these updated reference genomes and laid a foundation for using T2T-CHM13 and pangenome in follow-up epigenetics investigations.

## Methods

### Short-read sequencing DNA methylation data information

A total of 15 samples were collected because their corresponding biosamples were sequenced with at least three short-read sequencing technologies (Table S1). Specifically, MeDIP-seq, RRBS, and WGBS data sets for the GM12878 human B-lymphoblastoid cell line, H1 human embryonic stem cell (hESC) line, K562 human leukemia cell line, and H9 hESC line, as well as three MBD-seq samples for H9 hESCs, were retrieved from the Encyclopedia of DNA Elements (ENCODE) project and the NCBI Sequence Read Archive (SRA) database for calling CpGs^63,69^. In addition, 20 RRBS samples (tumor, *n* = 10; adjacent normal tissue, *n* = 10) were retrieved from GSE95654 for identification of colon cancer-associated CpGs^17^ (Table S4). 13 MeDIP-seq samples of East Asian ancestry (chronic lymphocytic leukemia, *n* = 8; normal B cell, *n* = 5) were downloaded from GSE136986 for determination of the benefit of the human pangenome reference in calling CpGs^54^ (Table S19).

### WGBS and RRBS data processing

WGBS and RRBS data were preprocessed according to the Quality Control, Trimming, and Alignment of Bisulfite-Seq Data Protocol^70^. First, quality control (QC) checks on the raw WGBS and RRBS data were applied using FastQC version 0.11.9 (https://www.bioinformatics.babraham.ac.uk/projects/fastqc/)^71^. To remove sequencing adapters and low-quality base reads, sequencing reads in WGBS and RRBS files were trimmed with Trim Galore version 0.6.6 (https://www.bioinformatics.babraham.ac.uk/projects/trim_galore/)^72^. These trimmed reads were then mapped to the human GRCh38.p13 (https://www.ncbi.nlm.nih.gov/assembly/GCF_000001405.39/) and T2T-CHM13 v2.0 (https://www.ncbi.nlm.nih.gov/assembly/GCF_009914755.1/) assemblies with Bismark version 0.22.3 (https://www.bioinformatics.babraham.ac.uk/projects/bismark/) using the default parameters^65^. In addition, the `deduplication` tool in Bismark was used to remove duplications in WGBS files. Next, non-CpG methylation reads in WGBS and RRBS files were removed using the Bismark filtration tool (options: `-s` for single-end file; and `-p` for paired-end file). Finally, the `bismark_methylation_extractor` (parameters: `--comprehensive --merge_non_CpG --bedGraph --counts`) and `coverage2cytosine` (parameters: `--merge_CpG`) modules in Bismark were used to extract cytosine methylation from the alignment reads.

### MBD-seq and MeDIP-seq data processing using T2T-CHM13

QC assessment of the sequencing reads from MBD-seq and MeDIP-seq data was performed using FastQC version 0.11.9 ^71^. Quality and adapter trimming were performed using Trim Galore version 0.6.6 ^72^. The trimmed reads were aligned to the human GRCh38.p13 and T2T-CHM13 v2.0 assemblies with Bowtie2 version 2.4.4 (https://github.com/BenLangmead/bowtie2; parameters: `-q --phred33 --sensitive -X 500 -p 2 -t`)^73^. SAMtools version 1.6 (https://github.com/samtools/samtools) was used to sort, filter (parameters: `-F 1804 -q 2`), and index aligned reads^74^. Polymerase chain reaction duplicate reads were marked and removed with Picard version 2.25.5 (http://broadinstitute.github.io/picard; parameters: `REMOVE_DUPLICATES=true ASSUME_SORTED=true`). The number of actual CpGs within sequencing reads from MBD-seq and MeDIP-seq samples was estimated by converting sequence alignments in BAM format into BED to obtain their corresponding genomic coordinates and then intersecting those genomic coordinates with all CpG coordinates of the reference genome using bedtools v2.30.0 with the bamtobed and intersect utilities^75^. The actual coordinates of all CpGs were obtained by grepping “CG” from the full genome sequences of GRCh38 and T2T-CHM13, separately.

### Detection of CpGs associated with colon cancer

To detect DNAm alterations in colon cancer, RRBS data from 10 tumors and 10 adjacent normal tissue samples were used. Totals of 1,245,858 and 1,165,567 CpGs in autosomes with coverage of at least 10× in each sample were aligned to T2T-CHM13 and GRCh38.p13, respectively. Differential DNAm levels between tumor and normal tissues were examined using the Mann– Whitney U test. Regarding multiple testing adjustments, permutation tests are commonly considered the gold standard in GWAS and are successfully applied to EWAS^76,77^. Therefore, FDR was estimated via permutation analysis by randomly sampling tumor/normal labels 1000 times and recomputing the *P*-values for DNAm differences for each permutation. After each permutation, we kept the most significant *P*-value per CpG and selected the *P*-value threshold that provided a 5% FDR (*P* = 3.25 × 10^−4^ for CpGs called by T2T-CHM13 and *P* = 2.06 × 10^−4^ for CpGs called by GRCh38). CpGs with DNAm differences > 0.10 and FDR < 0.05 were recognized as colon cancer-associated CpGs.

For identification of CpGs uniquely called using T2T-CHM13 compared with GRCh38 (i.e., additional CpGs), CpGs called by GRCh38.p13 were converted to T2T-CHM13 v2.0 using the UCSC liftOver tool (https://genome-store.ucsc.edu/) with the default parameters and the chain file generated by the T2T Consortium^1,78^. Genome coordinates for 1,164,472 CpGs were successfully converted to T2T-CHM13 v2.0. Intersections (*n* = 1,143,869) between CpGs called by the two reference genomes were identified using bedtools v2.30.0. Additional CpGs (*n* = 101,989) were defined by excluding all intersecting CpGs from the CpGs called using T2T-CHM13.

PCA was performed to obtain a comprehensive view of DNAm patterns of the 88 colon cancer-associated CpGs, which benefited from the additional CpGs called using T2T-CHM13, across samples using the *prcomp* function in the `stats` R package^79^. All statistical analyses in this study were performed using R (Version 4.0.3: www.r-project.org/)^79^.

### Enrichment analysis of target CpGs for known genomic elements

To calculate enrichment statistics for target CpGs presented in a specific genomic element, Pearson’s chi-squared test was used to determine statistical significance at the Benjamini– Hochberg FDR threshold of 0.05 ^80^. The UCSC Genome Browser (https://genome.ucsc.edu/cgi-bin/hgTables) was used to retrieve annotation files for the promoter regions (2 kb upstream), CpG island regions, and Comparative Annotation Toolkit (CAT) + Liftoff genes. The genomic coordinates for these annotation files were based on the human reference genome T2T-CHM13 v2.0.

### Pathway enrichment analysis of genes

We mapped genes for target CpGs to biological pathways to investigate their functional significance. The analysis was restricted to CpGs from promoter and gene body regions. Gene Ontology (GO) molecular functions, GO biological processes, GO cellular components, Reactome, and KEGG pathway analyses were performed using the g:Profiler toolset (https://biit.cs.ut.ee/gprofiler/gost)^81^. Pathways were deemed to be significantly enriched at a threshold of Benjamini–Hochberg FDR < 0.05.

### Identification of unambiguous CpGs using T2T-CHM13

In contrast to previous studies concerning cross-reactive probes using GRCh37/hg19 ^38–42,44,45^, we identified unambiguous (i.e., non–cross-reactive and non-mismatched) probes. As subsets of probes were poorly aligned to the more recent GRCh38 assembly, array probes could not be divided into two categories (cross-reactive and unambiguous probes) as was done with GRCh37/hg19, but were rather divided into three categories, i.e., cross-reactive (i.e., probes that match multiple regions), mismatched (i.e., probes that do not perfectly match the target region), and unambiguous (i.e., non–cross-reactive and non-mismatched probes uniquely mapping to the target region)^43^. “Mismatch” included the inability of some probes to map to GRCh38 and the extension bases of some probes obtained from alignments differed from the annotations in the manifest file of the array based on GRCh37/hg19 (e.g., type II probes extended to a non-guanine base)^43^. Cross-reactive and mismatched probes have been reported to have a greater deviation in DNAm differences between HM450K and WGBS and are therefore often masked^43^. Therefore, the set of unambiguous probes can be easily incorporated into epigenetics studies rather than removing numerous independent sets of probes with unique mapping issues, which would involve more operations.

To capture unambiguous probes, we mapped HM450K (*n* = 485,512), EPIC (*n* = 865,859), and EPICv2 (*n* = 936,190) probes to human GRCh38.p13 and T2T-CHM13 v2.0 reference genomes downloaded from the National Center for Biotechnology Information (NCBI). The sequences of array probes were extracted from the Illumina annotation files. For type II probe sequences containing R nucleotides, the R nucleotides were replaced with A or G. We used BLAT (BLAST-like alignment tool) to map probe sequences to four different types of genome for each reference genome: (1) an unmethylated bisulfite-treated genome with all Cs converted to Ts; (2) a methylated bisulfite-treated genome with only non-CpG Cs converted to Ts; (3) the reverse-complement of (1); and (4) the reverse-complement of (2)^82^. The following parameters were used to implement BLAT: `tileSize=11`, `stepSize=5`, and `repMatch=1,000,000`. For a probe to be considered unambiguous, it had to be: non-mismatched in the sequence alignment with at least 90% identity, at least 40 of 50 matching bases, no gaps, and the CpG locus had to be perfectly matched; and non–cross-reactive in the sequence alignment with unique hits aligned to the human genome^38,39^. As an example, a flowchart of unambiguous probe discovery in EPICv2 arrays using T2T-CHM13 is illustrated in Fig. S6. Ultimately, 430,719, 777,491, and 859,216 unambiguous probes were defined in the HM450K, EPIC, and EPICv2 arrays, respectively, using T2T-CHM13, while 434,633, 783,830, and 866,472, respectively, were defined using GRCh38.

### DNAm array data preprocessing

A single blood sample from a 92-year-old man who participated in this study was taken from the CRELES cohort, which is comprised of 2827 participants aged 60 or older from Costa Rica^83^. The EPIC array was used to measure methylation patterns of 6 replicates at a genome-wide level. The raw DNAm data were color corrected and background subtracted using GenomeStudio software. Data analysis was carried out using R version 4.0.3. Preprocessing was performed using the `minfi` and `methylumi` R packages^84,85^. To ensure effective quality control, the entire data set of 504 samples, including the 6 replicates, was assessed. A subset of probes was removed for QC purposes. These included SNP control probes, probes with a detection *P*-value > 0.05 in more than 1% of samples, probes with missing values in more than 5% of samples, probes with limited bead count < 3 in more than 5% of samples, and polymorphic CpG probes. Furthermore, samples were excluded if more than 1% of the probes were missing. A beta-mixture quantile (BMIQ) normalization method was implemented to account for differences between probe types I and II^86^. As whole-blood samples are cellularly heterogeneous, cell proportions were calculated using the Houseman method and then regressed against the DNAm data^87–89^.

The preprocessed and normalized data sets for the publicly available array samples used in this study were downloaded from the Gene Expression Omnibus (GEO) public repository with accessions GSE103505, GSE55763, and GSE43414, and additional QC was carried out^90^. The SNP control probes, probes with missing values in more than 5% of samples, and polymorphic CpG probes were removed from each data set. Moreover, samples were discarded if more than 1% of the probes were missing. Cell proportions in blood samples were estimated from the DNAm data and corrected using the Houseman method^87–89^. As the IMR-90 cell line consists of fibroblasts isolated from normal lung tissue, no further cell type composition correlation analyses were performed for these samples.

### Measurement of DNAm variability across technical replicates and DNAm differences between array and WGBS

The DNAm variability among technical replicates was calculated using the standard deviation. Although DNAm beta value is associated with standard deviation^91^, we did not place probes into mean methylation categories (i.e., hypo-, intermediate, and hypermethylation) as cross-reactivity can lead to mixtures of multiple signals and more intermediate methylation (Fig. S7). The DNAm change was determined as the absolute difference in DNAm value measurement between the arrays and WGBS. As HM450K and EPIC array probes were designed based on the GRCh37/hg19 reference genome assembly, to reduce the effect of varied assemblies, the hg19-based processed file for IMR-90 WGBS data that contained CpG coordinates and methylation levels was downloaded from GSE103505 ^92^. Only CpG sites that were accessible on both platforms and those with a minimum coverage depth of 10× with WGBS were included in the analysis. DNAm variability (and change) between subsets of CpGs compared using the Mann– Whitney U test was regarded as statistically significant at FDR < 0.05. The FDR was estimated from *P*-values using the Benjamini-Hochberg procedure^80^.

### Enrichment analysis of GRCh38-unique probes for repetitive regions

Annotation files for the repetitive elements of the human reference genome T2T-CHM13 v2.0 were retrieved from the UCSC Genome Browser (https://genome.ucsc.edu/cgi-bin/hgTables). Repetitive elements were grouped into four categories, as reported previously^2^: segmental duplications, satellites, LINEs/SINEs, and all others. LINE and SINE regions were merged, and overlaps with segmental duplications were removed. The regions overlapping both segmental duplication and LINE/SINE regions were removed from satellite regions. The other repetitive element types extracted from the RepeatMasker annotation that overlapped with the previous three repetitive element types were also removed. Fold enrichment was calculated as the ratio of observed values in GRCh38-unique probes to observed values in T2T-unique probes. Pearson’s chi-squared test was used to determine statistical significance at the FDR threshold of 0.05.

### Identification of differentially methylated CpGs (DMCs) in 24 cancer types

24 cancer types were used as examples in this analysis. There were obtained from the NCBI Gene Expression Omnibus (GEO; https://www.ncbi.nlm.nih.gov/geo/), The Cancer Genome Atlas (TCGA; https://tcgadata.nci.nih.gov), the International Cancer Genome Consortium (ICGC; https://dcc.icgc.org/), the TARGET Data Matrix (https://ocg.cancer.gov/programs/target/data-matrix), and the ArrayExpress databases (https://www.ebi.ac.uk/arrayexpress/)^90,93–95^, with an average of 350 samples (Table S15). To ensure the expansion of our findings, this study primarily used data from the HM450K. For EPIC data, only CpG probes that were shared between the HM450K and the EPIC array were used. The preprocessing of these DNAm array data was the same as shown for the publicly available samples mentioned above. ‘ComBat’ from the ‘sva’ R package was used to correct batch effects between datasets^96^. To identify DMCs in each cancer type, we used the Mann–Whitney U test to measure the DNAm differences between tumor and normal tissues. FDR values were computed using the Benjamini–Hochberg method^80^. CpGs with DNAm difference > 0.1 and FDR < 0.05 were considered statistically significant^97^.

### Short-read sequencing data processing using the human pangenome reference

QC assessment and adapter trimming of the sequencing reads were performed using FastQC version 0.11.9 ^71^ and Trim Galore version 0.6.6 ^72^ separately. The trimmed reads were aligned to the T2T-CHM13 reference genome graph and to the pangenome graph with `vg giraffe` (version v1.53.0; https://github.com/vgteam/vg/; parameters: `-t 14 -Z -m -d -o BAM`)^98–100^. The pangenome graph used here followed MC graph creation strategy^101^, as available from https://github.com/human-pangenomics/hpp_pangenome_resources^7^. To be noted, the MC graph included 44 of the 47 samples assembled (for evaluation purposes, HG002, HG005 and NA19240 were excluded in graph constructions) and added GRCh38 and CHM13 references to make the total number of haplotypes included 90 ^7^. The T2T-CHM13 reference genome graph was constructed by `vg construct` (parameters: `-C -R -t 1 -m 32`)^98^ using a VCF file of the 1000 genomes project recalled on T2T-CHM13 v2.0 (https://s3-us-west-2.amazonaws.com/human-pangenomics/index.html?prefix=T2T/CHM13/assemblies/variants/1000_Genomes_Project/chm13v2.0/unrelated_samples_2504/)^31^. SAMtools version 1.6 (https://github.com/samtools/samtools) was used to sort and generate consensus from aligned reads^74^. The number of actual CpGs within sequencing reads from MeDIP-seq samples was estimated by grepping “CG” from the consensus FASTA file.

### Identification of unambiguous CpGs using the human pangenome reference

As the human pangenome reference contains 94 haplotype-level assemblies, we mapped HM450K (*n* = 485,512), EPIC (*n* = 865,859), and EPICv2 (*n* = 936,190) probes to each of the haplotype-level assemblies separately using BLAT as described above. To obtain a pangenome-based unambiguous CpG set, CpGs determined to be unambiguous in no less than 95% of the 94 haplotype-level assemblies were chosen to reduce the effect of common variants. To obtain a population- or subpopulation-based unambiguous CpG set, CpGs determined to be unambiguous in no less than 95% of haplotype-level assemblies (or all of haplotype-level assemblies for the limited number (n < 5) of assemblies in populations and subpopulations) corresponding to the human population or subpopulation were chosen.

## Declarations

## Abbreviations

ACB: African Caribbean in Barbados

AFR: African

AMR: American

ASW: African Ancestry in Southwest US

BLAT: BLAST-like alignment tool

bp: base pair

ChIP-seq: chromatin immunoprecipitation sequencing

CHS: Chinese

CLM: Colombian in Medellin, Colombia

CpG: cytosine-phosphate-guanine dinucleotide

DMC: differentially methylated CpG

DNAm: DNA methylation

EAS: East Asian

ENCODE: Encyclopedia of DNA Elements

EPIC: Illumina HumanMethylationEPIC BeadChip

EPICv2: Infinium MethylationEPIC v2.0

EUR: European

ESN: Esan in Nigeria

EWAS: epigenome-wide association study

FDR: false discovery rate

GEO: NCBI Gene Expression Omnibus

GWD: Gambian in Western Division

HM450K: HumanMethylation450 BeadChip

HPRC: Human Pangenome Reference Consortium

KEGG: Kyoto encyclopedia of genes and genomes

KHV: Kinh in Ho Chi Minh City, Vietnam

MBD-seq: methylated DNA binding domain sequencing

MeDIP-seq: methylated DNA immunoprecipitation sequencing

MKK: Maasai in Kinyawa, Kenya

MSL: Mende in Sierra Leone

PC: principal component

PCA: principal component analysis

PEL: Peruvian in Lima, Peru

PUR: Puerto Rican in Puerto Rico

QC: quality control

RRBS: reduced representation bisulfite sequencing

SAS: South Asian

SNP: single nucleotide polymorphism

WGBS: whole-genome bisulfite sequencing

YRI: Yoruba in Ibadan

## Acknowledgements

We are grateful to Compute Canada’s National Systems for Scientific Computation for computational support. We are extremely grateful to our colleague, Alan Kerr, whose wonderful contribution to writing and style greatly improved the manuscript. We are grateful to the individuals who participated in the CRELES study for providing their personal information and biological specimens without payment or compensation, as well as to the Costa Rican institutions (CONAPAM, CCSS, INEC, TSE, and INISA) that collaborated in data and specimen collection. We are also grateful to the staff and fieldworkers at the Central American Population Center (CCP) of the University of Costa Rica (UCR) who made the CRELES study possible.

## Authors’ contributions

Z.D. conceived, designed, and performed the computational analyses, analyzed the data, and interpreted the results. J.L.M., D.H.R., and L.R.B. contributed DNA methylation data. J.W. and M.F. contributed ideas and participated in evaluating results and discussions. K.K. and M.S.K. supervised the study and M.S.K. obtained funding. Z.D. wrote the manuscript, with input from all authors. All authors approved the final version of the manuscript.

## Consent for publication

Not applicable.

## Availability of data and materials

The list of unambiguous probes (i.e., non–cross-reactive and non-mismatched probes uniquely mapping to the target region) on Illumina methylation arrays is available for download from https://github.com/functionalepigenomics/Illumina_Infinium_HumanMethylation_BeadChips_Annotation. The short-read sequencing and methylation array data used in this study are available from the NCBI Sequence Read Archive (SRA; https://www.ncbi.nlm.nih.gov/sra/) and the NCBI Gene Expression Omnibus (GEO; http://www.ncbi.nlm.nih.gov/geo/). The NCBI RefSeq genome database is available at https://ftp.ncbi.nlm.nih.gov/genomes/refseq. The CRELES DNA methylation data are not publicly available. Requests for restricted access to data can be submitted at http://www.creles.berkeley.edu/ following institutional review approval.

## Code availability

The scripts used to generate the figures and support the findings of this study are available in https://github.com/functionalepigenomics/Reference_Genome_Updates_on_Methylation_Studies.

### Ethics approval and consent to participate

The Committee on Ethics and Science (Comité Etico Científico) of the University of Costa Rica (UCR) granted approval for research in human subjects to CRELES (ref. VI-763-CEC-23-04). All participants provided written informed consent. The DNA samples used in this study, which are owned by the UCR, were transferred to the University of California Berkeley under a Non-Commercial Material Transfer Agreement and Notification of Transfer on July 3, 2013.

## Supplementary Information

### Additional File 1

**Table S1.** Characteristics of Short-Read Sequencing DNA Methylation Data

**Table S2.** Percentage of T2T-CHM13-unique CpGs found in sequences not included in GRCh38 across all T2T-CHM13-unique CpGs

**Table S3.** Number of Shared CpGs Across Samples for Each Short-Read Sequencing Method

**Table S4.** Characteristics of Colon Cancer RRBS Data

**Table S5.** Colon Cancer-Associated CpGs Identified From RRBS Samples

**Table S6.** Enrichment of Colon Cancer-Associated CpGs Unique to T2T-CHM13 in Comparison of Random CpG Sites in Genomic Elements

**Table S7.** Cancer Driver Genes Whose Promoters and Gene Bodies overlapped with 88 Additional Colon Cancer-Associated CpGs

**Table S8.** Cancer Driver Genes Whose DNA Methylation Alterations Were Uniquely Detected by 88 Additional Colon Cancer-Associated CpGs

**Table S9.** Pathway Enrichment for the 58 Genes with Promoters and Gene Bodies Overlapping 88 Additional Colon Cancer-Associated CpGs in Comparison With All Genes

**Table S10.** Pathway Enrichment for the 58 Genes with Promoters and Gene Bodies Overlapping 88 Additional Colon Cancer-Associated CpGs in Comparison With the 707 Genes with Promoters and Gene Bodies Overlapping 896 Colon Cancer-Associated CpGs Shared With GRCh38

**Table S11.** Comparison of T2T-CHM13-Defined Unambiguous Probes to Those in Previous Studies

**Table S12.** T2T-CHM13-Uniquely Defined Unambiguous Probes in HM450K and EPIC

**Table S13.** GRCh38-Uniquely Defined Unambiguous Probes in HM450K and EPIC

**Table S14.** Enrichment of GRCh38-Uniquely Defined Unambiguous Probes in Comparison With T2T-CHM13-Uniquely Defined Unambiguous Probes in Repetitive Regions

**Table S15.** Characteristics of Cancer Methylation Data Sets Used in This Study

**Table S16.** Enrichment of DMCs Uniquely Identified by T2T-CHM13 in Comparison of Array CpG Sites and DMCs Shared with GRCh38 in Genomic Elements

**Table S17.** Enrichment of T2T-Unique DMCs Compared to Array CpG Sites and GRCh38 Shared DMCs in Cancer Driver Genes

**Table S18.** Number of Pathways Enriched for Cancer Driver Genes Uniquely Overlapping T2T-Unique DMCs Compared to Genes Overlapping Array CpG Sites and GRCh38 Shared DMCs

**Table S19.** Percentage of Pangenome-unique CpGs found in sequences not included in T2T-CHM13 across all Pangenome-unique CpGs

**Table S20.** Enrichment of Pangenome-Unique DMCs Compared to Array CpG Sites and T2T-CHM13 Shared DMCs in Cancer Driver Genes

**Table S21.** Cancer Driver Genes with Promoters and Gene Bodies Overlapping EAS-Specific Unambiguous Probes from the HM450K Array

**Table S22.** Shared Probes Between HM450K and EPIC Arrays and Between EPIC and EPICv2 Arrays

### Additional File 2

Figure S1. Loadings using the first two principal components (PCs) for each RRBS sample resulting from PCA color-coded by disease state of colon cancer.

**Figure S2.** Colocalization of colon cancer-associated CpGs that benefited from the additional CpGs called using T2T-CHM13 (n = 88) with genomic elements. Some CpGs overlapped multiple genomic features and were counted for each feature independently.

**Figure S3.** DNAm variability of CpGs in HM450K (left) and EPIC (right) across one blood sample with six technical replicates in all CpG groups. Mean values are represented as circles in each box plot. Not significant (ns); FDR ≥ 0.05; * 0.01 ≤ FDR < 0.05; ** 0.001 ≤ FDR < 0.01; *** FDR < 0.001. Outliers are not shown in box plots.

**Figure S4.** DNAm variability of HM450K CpGs across 36 blood (left) and cerebellum (right) samples with two technical replicates each for all CpG groups. Mean values are represented as circles in each box plot. Not significant (ns); FDR ≥ 0.05; * 0.01 ≤ FDR < 0.05; ** 0.001 ≤ FDR < 0.01; *** FDR < 0.001. Outliers are not shown in box plots.

**Figure S5.** Boxplots illustrating the proportions of variance explained by the top two PCs for DMCs found only with T2T-CHM13 in various cancers. Each dot represents a type of cancer.

**Figure S6.** Flowchart of unambiguous probe discovery in the EPICv2 array using T2T-CHM13. Probes in the array were first aligned to the reference genome to determine whether they showed good alignment (i.e., sequence alignment with at least 90% identity, at least 40 of 50 matching bases, no gaps, and the CpG locus had to be perfectly matched); if not, they were classified as mismatched probes; if yes, they were further checked to determine whether they were unique alignments; if yes, they were classified as unambiguous probes (i.e., non–cross-reactive and non-mismatched probes uniquely mapping to the target region); otherwise, they were classified as cross-reactive probes.

**Figure S7.** Mean methylation density differences between ambiguous and unambiguous probes in HM450K (left, *n* = 4) and EPIC (right, *n* = 3) across technical replicates from the IMR-90 cell line. Ambiguous probes were defined as those with cross-reactivity and/or mismatch.

## References

1. Nurk, S., Koren, S., Rhie, A., Rautiainen, M., Bzikadze, A. V., Mikheenko, A., Vollger, M.R., Altemose, N., Uralsky, L., Gershman, A., et al. (2022). The complete sequence of a human genome. Science (80-.). 376, 44–53. 10.1126/science.abj6987.

2. Gershman, A., Sauria, M.E.G., Guitart, X., Vollger, M.R., Hook, P.W., Hoyt, S.J., Jain, M., Shumate, A., Razaghi, R., Koren, S., et al. (2022). Epigenetic patterns in a complete human genome. Science (80-.). 376, eabj5089. 10.1126/SCIENCE.ABJ5089/SUPPL_FILE/SCIENCE.ABJ5089_MDAR_REPRODUCIBILITY_CHECKLIST.PDF.

3. Greenberg, M.V.C., and Bourc’his, D. (2019). The diverse roles of DNA methylation in mammalian development and disease. Nat. Rev. Mol. Cell Biol. 20, 590–607. 10.1038/s41580-019-0159-6.

4. Van Dongen, J., Nivard, M.G., Willemsen, G., Hottenga, J.J., Helmer, Q., Dolan, C. V., Ehli, E.A., Davies, G.E., Van Iterson, M., Breeze, C.E., et al. (2016). Genetic and environmental influences interact with age and sex in shaping the human methylome. Nat. Commun. 7, 11115. 10.1038/ncomms11115.

5. Ebert, P., Audano, P.A., Zhu, Q., Rodriguez-Martin, B., Porubsky, D., Bonder, M.J., Sulovari, A., Ebler, J., Zhou, W., Mari, R.S., et al. (2021). Haplotype-resolved diverse human genomes and integrated analysis of structural variation. Science 372, eabf7117. 10.1126/SCIENCE.ABF7117.

6. Wang, T., Antonacci-Fulton, L., Howe, K., Lawson, H.A., Lucas, J.K., Phillippy, A.M., Popejoy, A.B., Asri, M., Carson, C., Chaisson, M.J.P., et al. (2022). The Human Pangenome Project: a global resource to map genomic diversity. Nature 604, 437–446. 10.1038/s41586-022-04601-8.

7. Liao, W.W., Asri, M., Ebler, J., Doerr, D., Haukness, M., Hickey, G., Lu, S., Lucas, J.K., Monlong, J., Abel, H.J., et al. (2023). A draft human pangenome reference. Nature 617, 312–324. 10.1038/s41586-023-05896-x.

8. Amarasinghe, S.L., Su, S., Dong, X., Zappia, L., Ritchie, M.E., and Gouil, Q. (2020). Opportunities and challenges in long-read sequencing data analysis. Genome Biol. 21, 1–16. 10.1186/S13059-020-1935-5.

9. Lister, R., Pelizzola, M., Dowen, R.H., Hawkins, R.D., Hon, G., Tonti-Filippini, J., Nery, J.R., Lee, L., Ye, Z., Ngo, Q.M., et al. (2009). Human DNA methylomes at base resolution show widespread epigenomic differences. Nature 462, 315–322. 10.1038/nature08514.

10. Meissner, A., Mikkelsen, T.S., Gu, H., Wernig, M., Hanna, J., Sivachenko, A., Zhang, X., Bernstein, B.E., Nusbaum, C., Jaffe, D.B., et al. (2008). Genome-scale DNA methylation maps of pluripotent and differentiated cells. Nature 454, 766–770. 10.1038/nature07107.

11. Serre, D., Lee, B.H., and Ting, A.H. (2010). MBD-isolated Genome Sequencing provides a high-throughput and comprehensive survey of DNA methylation in the human genome. Nucleic Acids Res. 38, 391–399. 10.1093/NAR/GKP992.

12. Jacinto, F. V., Ballestar, E., and Esteller, M. (2008). Methyl-DNA immunoprecipitation (MeDIP): hunting down the DNA methylome. Biotechniques 44, 35–43. 10.2144/000112708.

13. Vollger, M.R., Guitart, X., Dishuck, P.C., Mercuri, L., Harvey, W.T., Gershman, A., Diekhans, M., Sulovari, A., Munson, K.M., Lewis, A.P., et al. (2022). Segmental duplications and their variation in a complete human genome. Science (80-.). 376, eabj6965. 10.1126/SCIENCE.ABJ6965/SUPPL_FILE/SCIENCE.ABJ6965_MDAR_REPRODUCIBILITY_CHECKLIST.PDF.

14. Hoyt, S.J., Storer, J.M., Hartley, G.A., Grady, P.G.S., Gershman, A., de Lima, L.G., Limouse, C., Halabian, R., Wojenski, L., Rodriguez, M., et al. (2022). From telomere to telomere: The transcriptional and epigenetic state of human repeat elements. Science (80-.). 376. 10.1126/SCIENCE.ABK3112.

15. Altemose, N., Logsdon, G.A., Bzikadze, A. V., Sidhwani, P., Langley, S.A., Caldas, G. V., Hoyt, S.J., Uralsky, L., Ryabov, F.D., Shew, C.J., et al. (2022). Complete genomic and epigenetic maps of human centromeres. Science (80-.). 376, eabl4178. 10.1126/SCIENCE.ABL4178/SUPPL_FILE/SCIENCE.ABL4178_DATABASES_S1_TO_S21.ZIP.

16. Kumar, C.M., Ryan, D.P., and Langhorst, B.W. (2021). Correcting Methylation Calls in Clinically Relevant Low-Mappability Regions. bioRxiv. 10.1101/2021.10.04.463127.

17. Hanley, M.P., Hahn, M.A., Li, A.X., Wu, X., Lin, J., Wang, J., Choi, A.H., Ouyang, Z., Fong, Y., Pfeifer, G.P., et al. (2017). Genome-wide DNA methylation profiling reveals cancer-associated changes within early colonic neoplasia. Oncogene 36, 5035–5044. 10.1038/ONC.2017.130.

18. Raut, J.R., Guan, Z., Schrotz-King, P., and Brenner, H. (2020). Fecal DNA methylation markers for detecting stages of colorectal cancer and its precursors: A systematic review. Clin. Epigenetics 12, 122. 10.1186/s13148-020-00904-7.

19. Lenhard, B., Sandelin, A., and Carninci, P. (2012). Metazoan promoters: Emerging characteristics and insights into transcriptional regulation. Nat. Rev. Genet. 13, 233–245. 10.1038/nrg3163.

20. Deaton, A.M., Bird, A., and Deaton, M. (2011). CpG islands and the regulation of transcription CpG islands and the regulation of transcription. Genes Dev. 25, 1010–1022. 10.1101/gad.2037511.

21. Lou, S., Lee, H.M., Qin, H., Li, J.W., Gao, Z., Liu, X., Chan, L.L., Lam, V.K.L., So, W.Y., Wang, Y., et al. (2014). Whole-genome bisulfite sequencing of multiple individuals reveals complementary roles of promoter and gene body methylation in transcriptional regulation. Genome Biol. 15, 1–21. 10.1186/S13059-014-0408-0/FIGURES/11.

22. Irizarry, R.A., Ladd-Acosta, C., Wen, B., Wu, Z., Montano, C., Onyango, P., Cui, H., Gabo, K., Rongione, M., Webster, M., et al. (2009). The human colon cancer methylome shows similar hypo- and hypermethylation at conserved tissue-specific CpG island shores. Nat. Genet. 41, 178–186. 10.1038/ng.298.

23. Edgar, R., Tan, P.P.C., Portales-Casamar, E., and Pavlidis, P. (2014). Meta-analysis of human methylomes reveals stably methylated sequences surrounding CpG islands associated with high gene expression. Epigenetics and Chromatin 7, 1–13. 10.1186/1756-8935-7-28/COMMENTS.

24. Wu, H., Coskun, V., Tao, J., Xie, W., Ge, W., Yoshikawa, K., Li, E., Zhang, Y., and Sun, Y.E. (2010). Dnmt3a-Dependent Nonpromoter DNA Methylation Facilitates Transcription of Neurogenic Genes. Science 329, 444. 10.1126/SCIENCE.1190485.

25. Feinberg, A.P., Ohlsson, R., and Henikoff, S. (2006). The epigenetic progenitor origin of human cancer. Nat. Rev. Genet. 7, 21–33. 10.1038/nrg1748.

26. Liang, W.W., Lu, R.J.H., Jayasinghe, R.G., Foltz, S.M., Porta-Pardo, E., Geffen, Y., Wendl, M.C., Lazcano, R., Kolodziejczak, I., Song, Y., et al. (2023). Integrative multi-omic cancer profiling reveals DNA methylation patterns associated with therapeutic vulnerability and cell-of-origin. Cancer Cell 41, 1567–1585.e7. 10.1016/J.CCELL.2023.07.013/ATTACHMENT/A0A36515-D1FD-4BE3-AA64-7A355A6D49AC/MMC7.XLSX.

27. Martínez-Jiménez, F., Muiños, F., Sentís, I., Deu-Pons, J., Reyes-Salazar, I., Arnedo-Pac, C., Mularoni, L., Pich, O., Bonet, J., Kranas, H., et al. (2020). A compendium of mutational cancer driver genes. Nat. Rev. Cancer 20, 555–572. 10.1038/s41568-020-0290-x.

28. Liu, C.-H., Lai, Y.-L., Shen, P.-C., Liu, H.-C., Tsai, M.-H., Wang, Y.-D., Lin, W.-J., Chen, F.-H., Li, C.-Y., Wang, S.-C., et al. (2023). DriverDBv4: a multi-omics integration database for cancer driver gene research. Nucleic Acids Res. 2023, 1–7. 10.1093/NAR/GKAD1060.

29. Earley, Z.M., Lisicka, W., Sifakis, J.J., Aguirre-Gamboa, R., Kowalczyk, A., Barlow, J.T., Shaw, D.G., Discepolo, V., Tan, I.L., Gona, S., et al. (2023). GATA4 controls regionalization of tissue immunity and commensal-driven immunopathology. Immunity 56, 43–57. 10.1016/j.immuni.2022.12.009.

30. Buikhuisen, J.Y., Barila, P.M.G., Torang, A., Dekker, D., de Jong, J.H., Cameron, K., Vitale, S., Stassi, G., van Hooff, S.R., Castro, M.A.A., et al. (2021). AKT3 Expression in Mesenchymal Colorectal Cancer Cells Drives Growth and Is Associated with Epithelial-Mesenchymal Transition. Cancers (Basel). 13, 1–19. 10.3390/CANCERS13040801.

31. Aganezov, S., Yan, S.M., Soto, D.C., Kirsche, M., Zarate, S., Avdeyev, P., Taylor, D.J., Shafin, K., Shumate, A., Xiao, C., et al. (2022). A complete reference genome improves analysis of human genetic variation. Science 376, eabl3533. 10.1126/SCIENCE.ABL3533.

32. Fang, J.Y., and Richardson, B.C. (2005). The MAPK signalling pathways and colorectal cancer. Lancet Oncol. 6, 322–327. 10.1016/S1470-2045(05)70168-6.

33. Engelhardt, S. (2007). Alternative signaling: cardiomyocyte β1-adrenergic receptors signal through EGFRs. J. Clin. Invest. 117, 2396. 10.1172/JCI33135.

34. Li, N., Xi, Y., Tinsley, H.N., Gurpinar, E., Gary, B.D., Zhu, B., Li, Y., Chen, X., Keeton, A.B., Abadi, A.H., et al. (2013). Sulindac selectively inhibits colon tumor cell growth by activating the cGMP/PKG pathway to suppress Wnt/β-catenin signaling. Mol. Cancer Ther. 12, 1848. 10.1158/1535-7163.MCT-13-0048.

35. Hanahan, D., and Weinberg, R.A. (2011). Hallmarks of cancer: The next generation. Cell 144, 646–674. 10.1016/J.CELL.2011.02.013/ATTACHMENT/3F528E16-8B3C-4D8D-8DE5-43E0C98D8475/MMC1.PDF.

36. Hanahan, D. (2022). Hallmarks of Cancer: New Dimensions. Cancer Discov. 12, 31–46. 10.1158/2159-8290.CD-21-1059.

37. Hop, P.J., Zwamborn, R.A.J., Hannon, E.J., Dekker, A.M., van Eijk, K.R., Walker, E.M., Iacoangeli, A., Jones, A.R., Shatunov, A., Al Khleifat, A., et al. (2020). Cross-reactive probes on Illumina DNA methylation arrays: a large study on ALS shows that a cautionary approach is warranted in interpreting epigenome-wide association studies. NAR genomics Bioinforma. 2, lqaa105. 10.1093/NARGAB/LQAA105.

38. Chen, Y. an, Choufani, S., Ferreira, J.C., Grafodatskaya, D., Butcher, D.T., and Weksberg, R. (2011). Sequence overlap between autosomal and sex-linked probes on the Illumina HumanMethylation27 microarray. Genomics 97, 214–222. 10.1016/J.YGENO.2010.12.004.

39. Price, M.E., Cotton, A.M., Lam, L.L., Farré, P., Emberly, E., Brown, C.J., Robinson, W.P., and Kobor, M.S. (2013). Additional annotation enhances potential for biologically-relevant analysis of the Illumina Infinium HumanMethylation450 BeadChip array. Epigenetics and Chromatin 6, 4. 10.1186/1756-8935-6-4.

40. Naeem, H., Wong, N.C., Chatterton, Z., Hong, M.K.H., Pedersen, J.S., Corcoran, N.M., Hovens, C.M., and Macintyre, G. (2014). Reducing the risk of false discovery enabling identification of biologically significant genome-wide methylation status using the HumanMethylation450 array. BMC Genomics 15, 1–15. 10.1186/1471-2164-15-51/FIGURES/4.

41. Chen, Y.A., Lemire, M., Choufani, S., Butcher, D.T., Grafodatskaya, D., Zanke, B.W., Gallinger, S., Hudson, T.J., and Weksberg, R. (2013). Discovery of cross-reactive probes and polymorphic CpGs in the Illumina Infinium HumanMethylation450 microarray. Epigenetics 8, 203. 10.4161/EPI.23470.

42. Pidsley, R., Zotenko, E., Peters, T.J., Lawrence, M.G., Risbridger, G.P., Molloy, P., Van Djik, S., Muhlhausler, B., Stirzaker, C., and Clark, S.J. (2016). Critical evaluation of the Illumina MethylationEPIC BeadChip microarray for whole-genome DNA methylation profiling. Genome Biol. 17, 1–17. 10.1186/S13059-016-1066-1/FIGURES/6.

43. Zhou, W., Laird, P.W., and Shen, H. (2017). Comprehensive characterization, annotation and innovative use of Infinium DNA methylation BeadChip probes. Nucleic Acids Res. 45, e22. 10.1093/NAR/GKW967.

44. McCartney, D.L., Walker, R.M., Morris, S.W., McIntosh, A.M., Porteous, D.J., and Evans, K.L. (2016). Identification of polymorphic and off-target probe binding sites on the Illumina Infinium MethylationEPIC BeadChip. Genomics Data 9, 22–24. 10.1016/J.GDATA.2016.05.012.

45. Benton, M.C., Johnstone, A., Eccles, D., Harmon, B., Hayes, M.T., Lea, R.A., Griffiths, L., Hoffman, E.P., Stubbs, R.S., and Macartney-Coxson, D. (2015). An analysis of DNA methylation in human adipose tissue reveals differential modification of obesity genes before and after gastric bypass and weight loss. Genome Biol. 16, 1–21. 10.1186/S13059-014-0569-X/TABLES/6.

46. Li, M., Deng, X., Zhou, D., Liu, X., Dai, J., and Liu, Q. (2024). A Novel Methylation-based Model for Prognostic Prediction in Lung Adenocarcinoma. Curr. Genomics 25, 26. 10.2174/0113892029277397231228062412.

47. Wang, X., Zhang, L., Liang, Q., Wong, C.C., Chen, H., Gou, H., Dong, Y., Liu, W., Li, Z., Ji, J., et al. (2022). DUSP5P1 promotes gastric cancer metastasis and platinum drug resistance. Oncogenesis 11, 1–11. 10.1038/s41389-022-00441-3.

48. Ghafouri-Fard, S., Khoshbakht, T., Hussen, B.M., Taheri, M., and Mokhtari, M. (2022). A review on the role of LINC01133 in cancers. Cancer Cell Int. 22, 1–10. 10.1186/S12935-022-02690-Z/TABLES/3.

49. Vega-Benedetti, A.F., Loi, E., Moi, L., Blois, S., Fadda, A., Antonelli, M., Arcella, A., Badiali, M., Giangaspero, F., Morra, I., et al. (2019). Clustered protocadherins methylation alterations in cancer. Clin. Epigenetics 11, 1–20. 10.1186/S13148-019-0695-0/TABLES/5.

50. Zhu, H., Blake, S., Chan, K.T., Pearson, R.B., and Kang, J. (2018). Cystathionine β-Synthase in Physiology and Cancer. Biomed Res. Int. 2018, 3205125. 10.1155/2018/3205125.

51. Zhang, Y.L., Wang, R.C., Cheng, K., Ring, B.Z., and Su, L. (2017). Roles of Rap1 signaling in tumor cell migration and invasion. Cancer Biol. Med. 14, 90. 10.20892/J.ISSN.2095-3941.2016.0086.

52. Monteith, G.R., Prevarskaya, N., and Roberts-Thomson, S.J. (2017). The calcium–cancer signalling nexus. Nat. Rev. Cancer 17, 373–380. 10.1038/nrc.2017.18.

53. Hoxhaj, G., and Manning, B.D. (2019). The PI3K–AKT network at the interface of oncogenic signalling and cancer metabolism. Nat. Rev. Cancer 20, 74–88. 10.1038/s41568-019-0216-7.

54. Kim, M., Lee, E., Zang, D.Y., Kim, H.J., Kim, H.Y., Han, B., Kim, H.S., Kang, H.J., Hwang, S., and Lee, Y.K. (2020). Novel genes exhibiting DNA methylation alterations in Korean patients with chronic lymphocytic leukaemia: a methyl-CpG-binding domain sequencing study. Sci. Rep. 10, 1085. 10.1038/S41598-020-57919-6.

55. Lappalainen, I., Lopez, J., Skipper, L., Hefferon, T., Spalding, J.D., Garner, J., Chen, C., Maguire, M., Corbett, M., Zhou, G., et al. (2013). dbVar and DGVa: public archives for genomic structural variation. Nucleic Acids Res. 41, D936. 10.1093/NAR/GKS1213.

56. Hu, S., Tao, J., Peng, M., Ye, Z., Chen, Z., Chen, H., Yu, H., Wang, B., Fan, J.B., and Ni, B. (2023). Accurate detection of early-stage lung cancer using a panel of circulating cell-free DNA methylation biomarkers. Biomark. Res. 11, 45. 10.1186/S40364-023-00486-5.

57. Hurley, C.K., Kempenich, J., Wadsworth, K., Sauter, J., Hofmann, J.A., Schefzyk, D., Schmidt, A.H., Galarza, P., Cardozo, M.B.R., Dudkiewicz, M., et al. (2020). Common, intermediate and well-documented HLA alleles in world populations: CIWD version 3.0.0. HLA 95, 516–531. 10.1111/TAN.13811.

58. Novembre, J., Johnson, T., Bryc, K., Kutalik, Z., Boyko, A.R., Auton, A., Indap, A., King, K.S., Bergmann, S., Nelson, M.R., et al. (2008). Genes mirror geography within Europe. Nature 456, 98–101. 10.1038/NATURE07331.

59. Campbell, M.C., and Tishkoff, S.A. (2008). African genetic diversity: implications for human demographic history, modern human origins, and complex disease mapping. Annu. Rev. Genomics Hum. Genet. 9, 403–433. 10.1146/ANNUREV.GENOM.9.081307.164258.

60. Logsdon, G.A., Vollger, M.R., and Eichler, E.E. (2020). Long-read human genome sequencing and its applications. Nat. Rev. Genet. 21, 597–614. 10.1038/s41576-020-0236-x.

61. Stunnenberg, H.G., Abrignani, S., Adams, D., de Almeida, M., Altucci, L., Amin, V., Amit, I., Antonarakis, S.E., Aparicio, S., Arima, T., et al. (2016). The International Human Epigenome Consortium: A Blueprint for Scientific Collaboration and Discovery. Cell 167, 1145–1149. 10.1016/J.CELL.2016.11.007.

62. Bernstein, B.E., Stamatoyannopoulos, J.A., Costello, J.F., Ren, B., Milosavljevic, A., Meissner, A., Kellis, M., Marra, M.A., Beaudet, A.L., Ecker, J.R., et al. (2010). The NIH Roadmap Epigenomics Mapping Consortium. Nat. Biotechnol. 28, 1045. 10.1038/NBT1010-1045.

63. Dunham, I., Kundaje, A., Aldred, S.F., Collins, P.J., Davis, C.A., Doyle, F., Epstein, C.B., Frietze, S., Harrow, J., Kaul, R., et al. (2012). An integrated encyclopedia of DNA elements in the human genome. Nature 489, 57–74. 10.1038/nature11247.

64. Pappalardo, X.G., and Barra, V. (2021). Losing DNA methylation at repetitive elements and breaking bad. Epigenetics Chromatin 2021 141 14, 1–21. 10.1186/S13072-021-00400-Z.

65. Krueger, F., and Andrews, S.R. (2011). Bismark: A flexible aligner and methylation caller for Bisulfite-Seq applications. Bioinformatics 27, 1571–1572. 10.1093/bioinformatics/btr167.

66. Xi, Y., and Li, W. (2009). BSMAP: Whole genome bisulfite sequence MAPping program. BMC Bioinformatics 10, 1–9. 10.1186/1471-2105-10-232/COMMENTS.

67. Hernández, H.G., Tse, M.Y., Pang, S.C., Arboleda, H., and Forero, D.A. (2013). Optimizing methodologies for PCR-based DNA methylation analysis. Biotechniques 55, 181–197. 10.2144/000114087/ASSET/IMAGES/LARGE/TABLE5.JPEG.

68. Kurdyukov, S., and Bullock, M. (2016). DNA Methylation Analysis: Choosing the Right Method. Biology (Basel). 5, 3. 10.3390/biology5010003.

69. Leinonen, R., Sugawara, H., and Shumway, M. (2011). The Sequence Read Archive. Nucleic Acids Res. 39, D19. 10.1093/NAR/GKQ1019.

70. Krueger, F., Kreck, B., Franke, A., and Andrews, S.R. (2012). DNA methylome analysis using short bisulfite sequencing data. Nat. Methods 9, 145–151. 10.1038/NMETH.1828.

71. Andrews, S. (2015). FASTQC a quality control tool for high throughput sequence data. https://www.bioinformatics.babraham.ac.uk/projects/fastqc/.

72. Krueger, F. (2016). Trim Galore. Babraham Bioinforma. http://www.bioinformatics.babraham.ac.uk/projects/trimgalore/.

73. Langmead, B., and Salzberg, S.L. (2012). Fast gapped-read alignment with Bowtie 2. Nat. Methods 9, 357. 10.1038/NMETH.1923.

74. Danecek, P., Bonfield, J.K., Liddle, J., Marshall, J., Ohan, V., Pollard, M.O., Whitwham, A., Keane, T., McCarthy, S.A., Davies, R.M., et al. (2021). Twelve years of SAMtools and BCFtools. Gigascience 10, giab008. 10.1093/GIGASCIENCE/GIAB008.

75. Quinlan, A.R., and Hall, I.M. (2010). BEDTools: a flexible suite of utilities for comparing genomic features. Bioinformatics 26, 841–842. 10.1093/BIOINFORMATICS/BTQ033.

76. Gao, X. (2011). Multiple testing corrections for imputed SNPs. Genet. Epidemiol. 35, 154. 10.1002/GEPI.20563.

77. Saffari, A., Silver, M.J., Zavattari, P., Moi, L., Columbano, A., Meaburn, E.L., and Dudbridge, F. (2018). Estimation of a significance threshold for epigenome wide association studies. Genet. Epidemiol. 42, 20. 10.1002/GEPI.22086.

78. Hinrichs, A.S., Karolchik, D., Baertsch, R., Barber, G.P., Bejerano, G., Clawson, H., Diekhans, M., Furey, T.S., Harte, R.A., Hsu, F., et al. (2006). The UCSC Genome Browser Database: update 2006. Nucleic Acids Res. 34, D590–8. 10.1093/NAR/GKJ144.

79. Team, R.C. (2021). R: A language and environment for statistical computing. R Foundation for Statistical Computing, Vienna, Austria. 2012. 27.

80. Benjamini, Y., and Hochberg, Y. (1995). Controlling the false discovery rate: a practical and powerful approach to multiple testing. J. R. Stat. Soc. Ser. B 57, 289–300. 10.1111/j.2517-6161.1995.tb02031.x.

81. Raudvere, U., Kolberg, L., Kuzmin, I., Arak, T., Adler, P., Peterson, H., and Vilo, J. (2019). g:Profiler: a web server for functional enrichment analysis and conversions of gene lists (2019 update). Nucleic Acids Res. 47, W191–W198. 10.1093/NAR/GKZ369.

82. Kent, W.J. (2002). BLAT - The BLAST-like alignment tool. Genome Res. 12, 656–664. 10.1101/gr.229202. Article published online before March 2002.

83. Brenes-Camacho, G., and Rosero-Bixby, L. (2009). Differentials by socioeconomic status and institutional characteristics in preventive service utilization by older persons in Costa Rica. J. Aging Health 21, 730–758. 10.1177/0898264309338299.

84. Aryee, M.J., Jaffe, A.E., Corrada-Bravo, H., Ladd-Acosta, C., Feinberg, A.P., Hansen, K.D., and Irizarry, R.A. (2014). Minfi: a flexible and comprehensive bioconductor package for the analysis of Infinium DNA methylation microarrays. Bioinformatics 30, 1363–1369. 10.1093/bioinformatics/btu049.

85. Davis, S., Du, P., Bilke, S., Triche, T., and Bootwalla, O. (2012). Methylumi: handle Illumina methylation data.

86. Teschendorff, A.E., Marabita, F., Lechner, M., Bartlett, T., Tegner, J., Gomez-Cabrero, D., and Beck, S. (2013). A beta-mixture quantile normalization method for correcting probe design bias in Illumina Infinium 450 k DNA methylation data. Bioinformatics 29, 189–196. 10.1093/bioinformatics/bts680.

87. Bakulski, K.M., Feinberg, J.I., Andrews, S. V., Yang, J., Brown, S., L. McKenney, S., Witter, F., Walston, J., Feinberg, A.P., and Fallin, M.D. (2016). DNA methylation of cord blood cell types: applications for mixed cell birth studies. Epigenetics 11, 354–362. 10.1080/15592294.2016.1161875.

88. Jaffe, A.E., and Irizarry, R.A. (2014). Accounting for cellular heterogeneity is critical in epigenome-wide association studies. Genome Biol. 15, R31. 10.1186/gb-2014-15-2-r31.

89. Houseman, E.A., Accomando, W.P., Koestler, D.C., Christensen, B.C., Marsit, C.J., Nelson, H.H., Wiencke, J.K., and Kelsey, K.T. (2012). DNA methylation arrays as surrogate measures of cell mixture distribution. BMC Bioinformatics 13, 86. 10.1186/1471-2105-13-86.

90. Barrett, T., Wilhite, S.E., Ledoux, P., Evangelista, C., Kim, I.F., Tomashevsky, M., Marshall, K.A., Phillippy, K.H., Sherman, P.M., Holko, M., et al. (2013). NCBI GEO: archive for functional genomics data sets--update. Nucleic Acids Res. 41, D991–5. 10.1093/NAR/GKS1193.

91. Xu, Z., and Taylor, J.A. (2021). Reliability of DNA methylation measures using Illumina methylation BeadChip. Epigenetics 16, 495–502. 10.1080/15592294.2020.1805692.

92. Suzuki, M., Liao, W., Wos, F., Johnston, A.D., DeGrazia, J., Ishii, J., Bloom, T., Zody, M.C., Germer, S., and Greally, J.M. (2018). Whole-genome bisulfite sequencing with improved accuracy and cost. Genome Res. 28, 1364–1371. 10.1101/GR.232587.117.

93. Weinstein, J.N., Collisson, E.A., Mills, G.B., Shaw, K.R.M., Ozenberger, B.A., Ellrott, K., Sander, C., Stuart, J.M., Chang, K., Creighton, C.J., et al. (2013). The cancer genome atlas pan-cancer analysis project. Nat. Genet. 45, 1113–1120. 10.1038/ng.2764.

94. Athar, A., Füllgrabe, A., George, N., Iqbal, H., Huerta, L., Ali, A., Snow, C., Fonseca, N.A., Petryszak, R., Papatheodorou, I., et al. (2019). ArrayExpress update - from bulk to single-cell expression data. Nucleic Acids Res. 47, D711–D715. 10.1093/NAR/GKY964.

95. Zhang, J., Bajari, R., Andric, D., Gerthoffert, F., Lepsa, A., Nahal-Bose, H., Stein, L.D., and Ferretti, V. (2019). The international cancer genome consortium data portal. Nat. Biotechnol. 37, 367–369. 10.1038/s41587-019-0055-9.

96. Leek, J.T., Johnson, W.E., Parker, H.S., Jaffe, A.E., and Storey, J.D. (2012). The sva package for removing batch effects and other unwanted variation in high-throughput experiments. Bioinformatics 28, 882. 10.1093/BIOINFORMATICS/BTS034.

97. Dong, Z., and Zhou, H. (2022). Pan-cancer landscape of aberrant DNA Methylation across childhood Cancers: Molecular Characteristics and Clinical relevance. Exp. Hematol. Oncol. 11, 1–5. 10.1186/S40164-022-00339-1/FIGURES/2.

98. Garrison, E., Sirén, J., Novak, A.M., Hickey, G., Eizenga, J.M., Dawson, E.T., Jones, W., Garg, S., Markello, C., Lin, M.F., et al. (2018). Variation graph toolkit improves read mapping by representing genetic variation in the reference. Nat. Biotechnol. 36, 875–879. 10.1038/nbt.4227.

99. Sirén, J., Monlong, J., Chang, X., Novak, A.M., Eizenga, J.M., Markello, C., Sibbesen, J.A., Hickey, G., Chang, P.C., Carroll, A., et al. (2021). Pangenomics enables genotyping of known structural variants in 5202 diverse genomes. Science (80-.). 374, abg8871. 10.1126/SCIENCE.ABG8871/SUPPL_FILE/SCIENCE.ABG8871_MDAR_REPRODUCIBILITY_CHECKLIST.PDF.

100. Sirén, J., and Paten, B. (2022). GBZ file format for pangenome graphs. Bioinformatics 38, 5012–5018. 10.1093/BIOINFORMATICS/BTAC656.

101. Hickey, G., Monlong, J., Ebler, J., Novak, A.M., Eizenga, J.M., Gao, Y., Abel, H.J., Antonacci-Fulton, L.L., Asri, M., Baid, G., et al. (2023). Pangenome graph construction from genome alignments with Minigraph-Cactus. Nat. Biotechnol. 42, 663–673. 10.1038/s41587-023-01793-w.

