## Supplementary Figures for "Complete Reference Genome and Pangenome Expand Biologically Relevant Information for Genome-Wide DNA Methylation Analysis Using Short-Read Sequencing and Array Data"

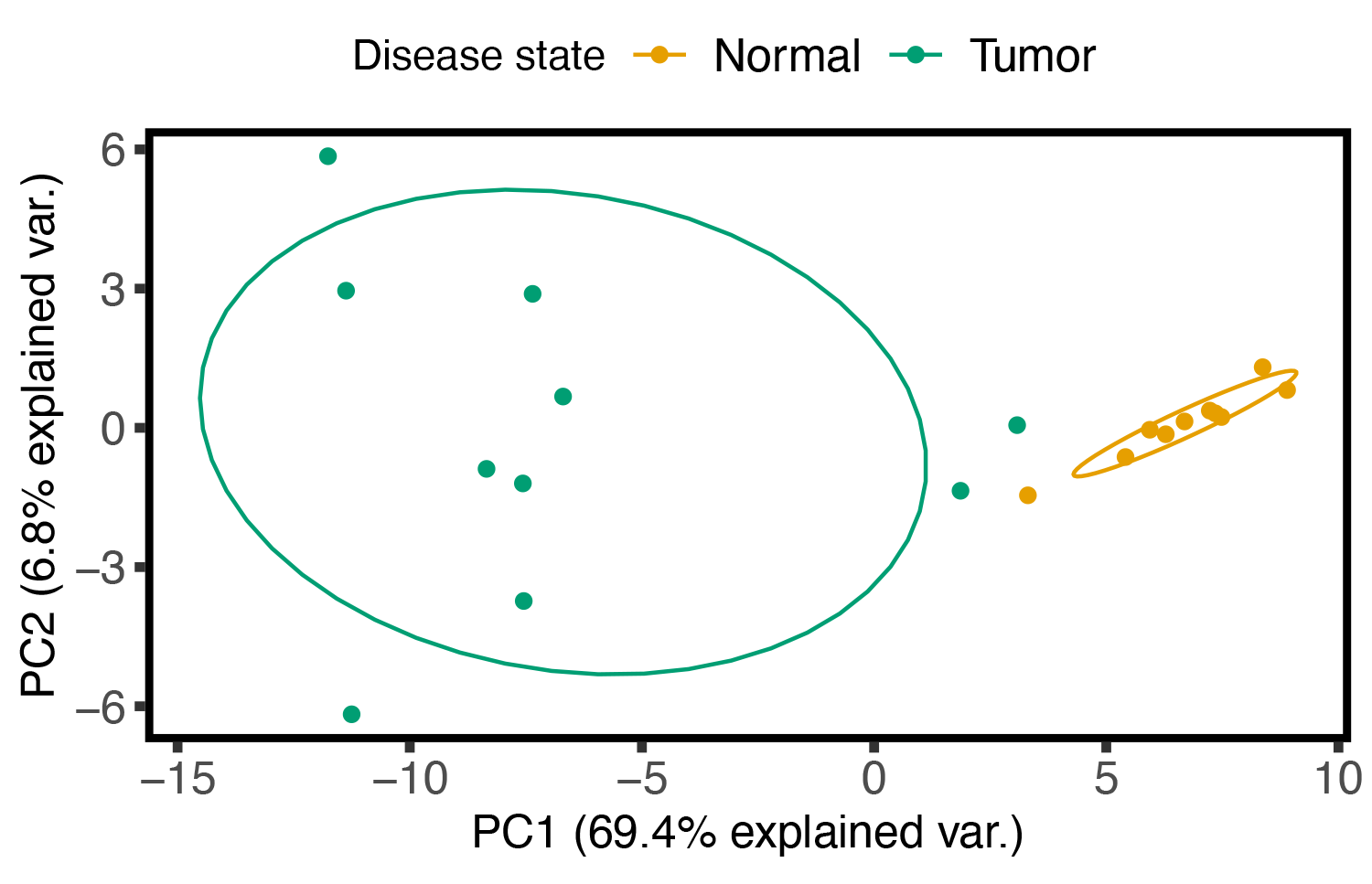

**Figure S1. Loadings using the first two principal components (PCs) for each RRBS sample resulting from PCA color-coded by disease state of colon cancer.**

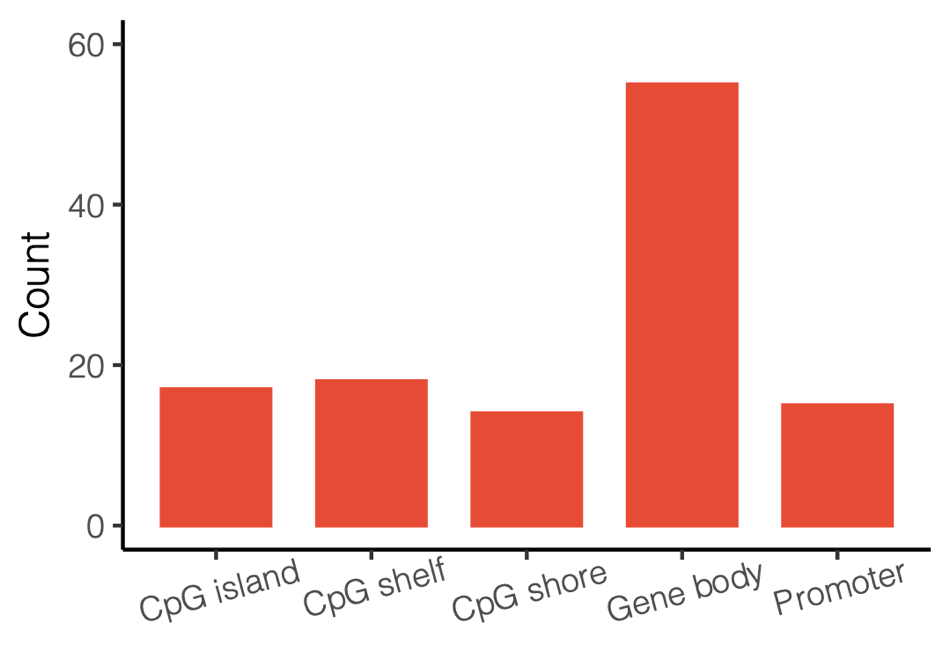

**Figure S2. Colocalization of colon cancer-associated CpGs that benefited from the additional CpGs called using T2T-CHM13 (n = 88) with genomic elements.** Some CpGs overlapped multiple genomic features and were counted for each feature independently.

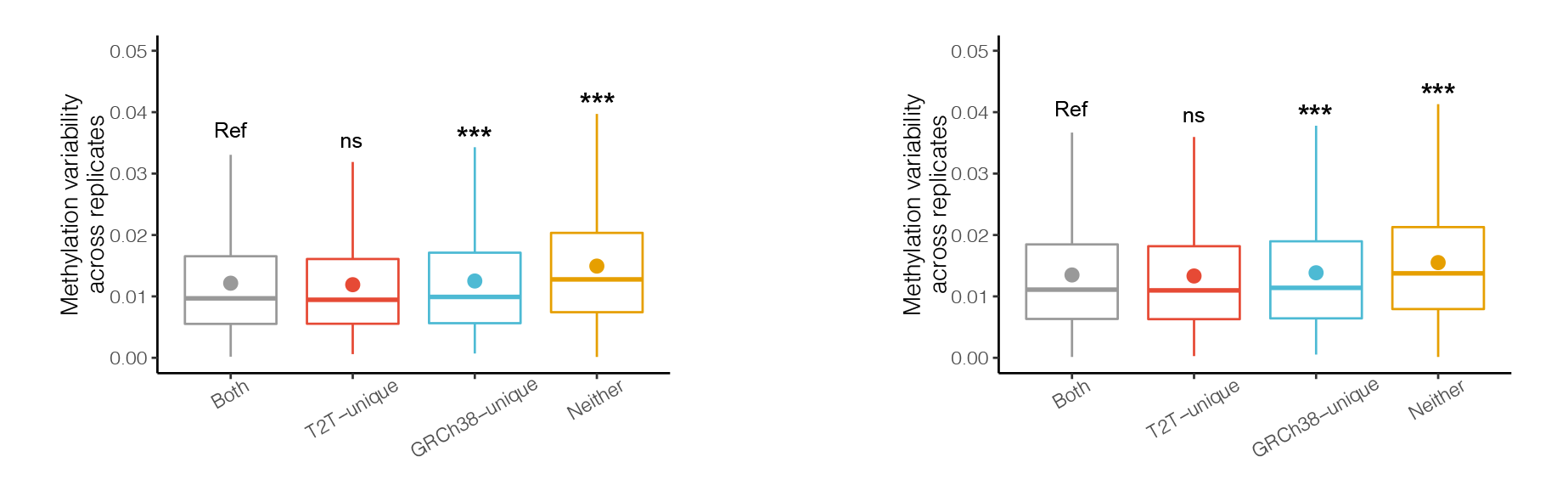

**Figure S3. DNAm variability of CpGs in HM450K (left) and EPIC (right) across one blood sample with six technical replicates in all CpG groups.** Mean values are represented as circles in each box plot. Not significant (ns); FDR ≥ 0.05; * 0.01 ≤ FDR < 0.05; ** 0.001 ≤ FDR < 0.01; *** FDR < 0.001. Outliers are not shown in box plots.

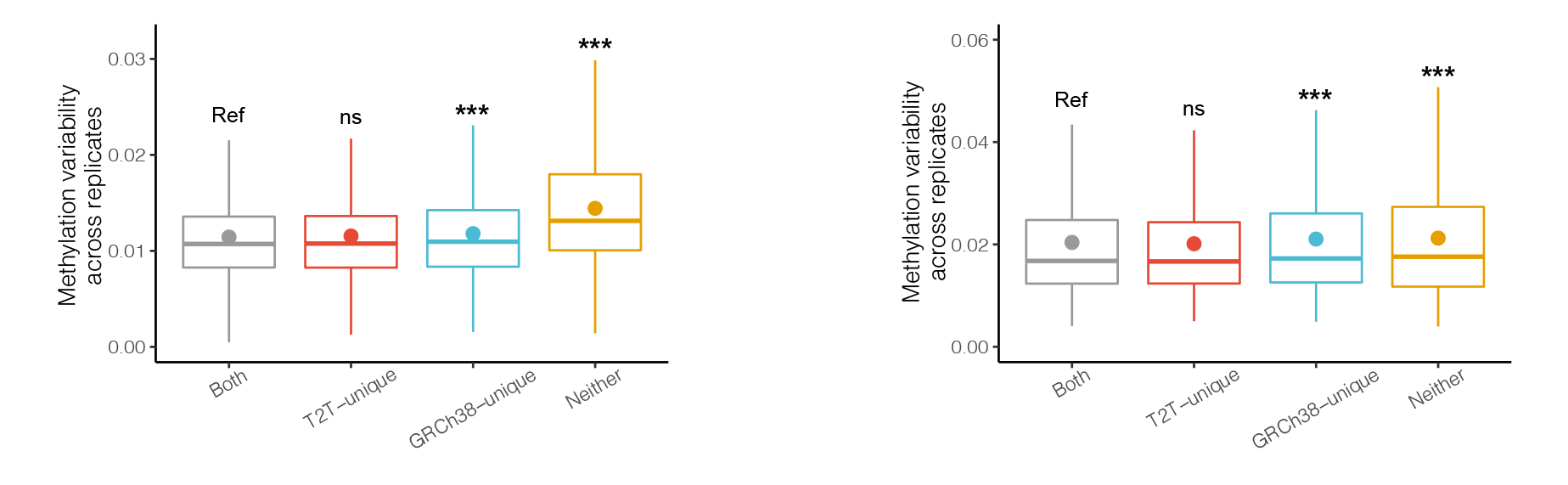

**Figure S4. DNAm variability of HM450K CpGs across 36 blood (left) and cerebellum (right) samples with two technical replicates each for all CpG groups.** Mean values are represented as circles in each box plot. Not significant (ns); FDR ≥ 0.05; * 0.01 ≤ FDR < 0.05; ** 0.001 ≤ FDR < 0.01; *** FDR < 0.001. Outliers are not shown in box plots.

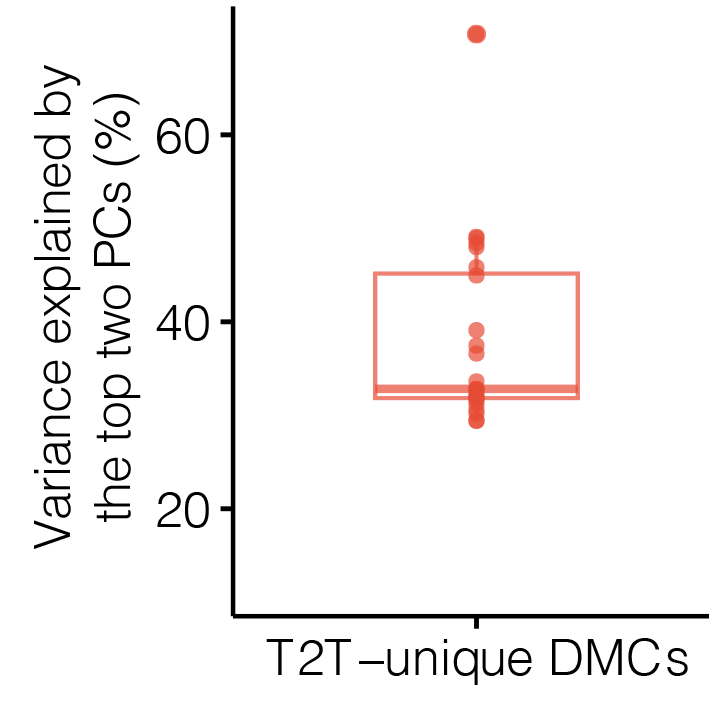

**Figure S5. Boxplots illustrating the proportions of variance explained by the top two PCs for DMCs found only with T2T-CHM13 in various cancers.** Each dot represents a type of cancer.

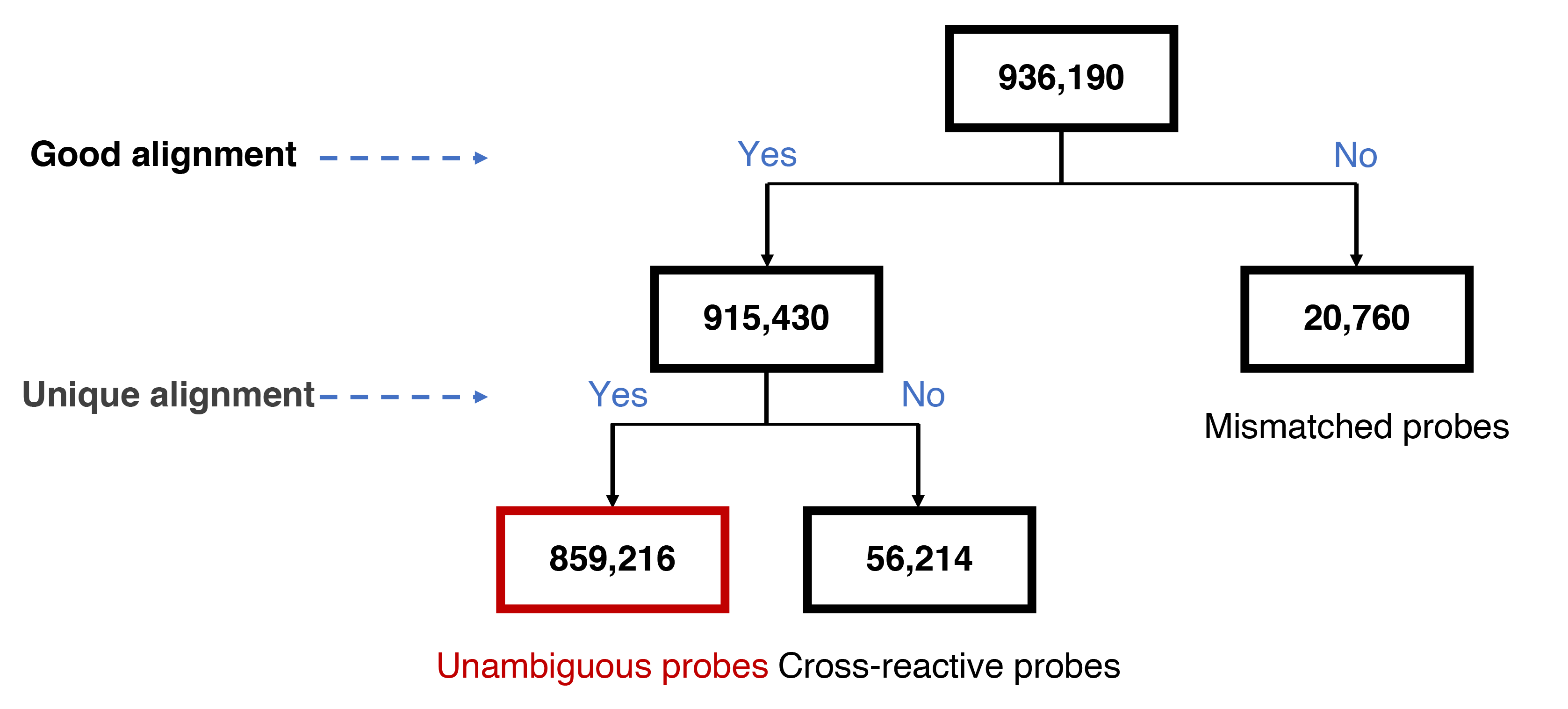

**Figure S6. Flowchart of unambiguous probe discovery in the EPICv2 array using T2T-CHM13.** Probes in the array were first aligned to the reference genome to determine whether they showed good alignment (i.e., sequence alignment with at least 90% identity, at least 40 of 50 matching bases, no gaps, and the CpG locus had to be perfectly matched); if not, they were classified as mismatched probes; if yes, they were further checked to determine whether they were unique alignments; if yes, they were classified as unambiguous probes (i.e., non–cross-reactive and non-mismatched probes uniquely mapping to the target region); otherwise, they were classified as cross-reactive probes.

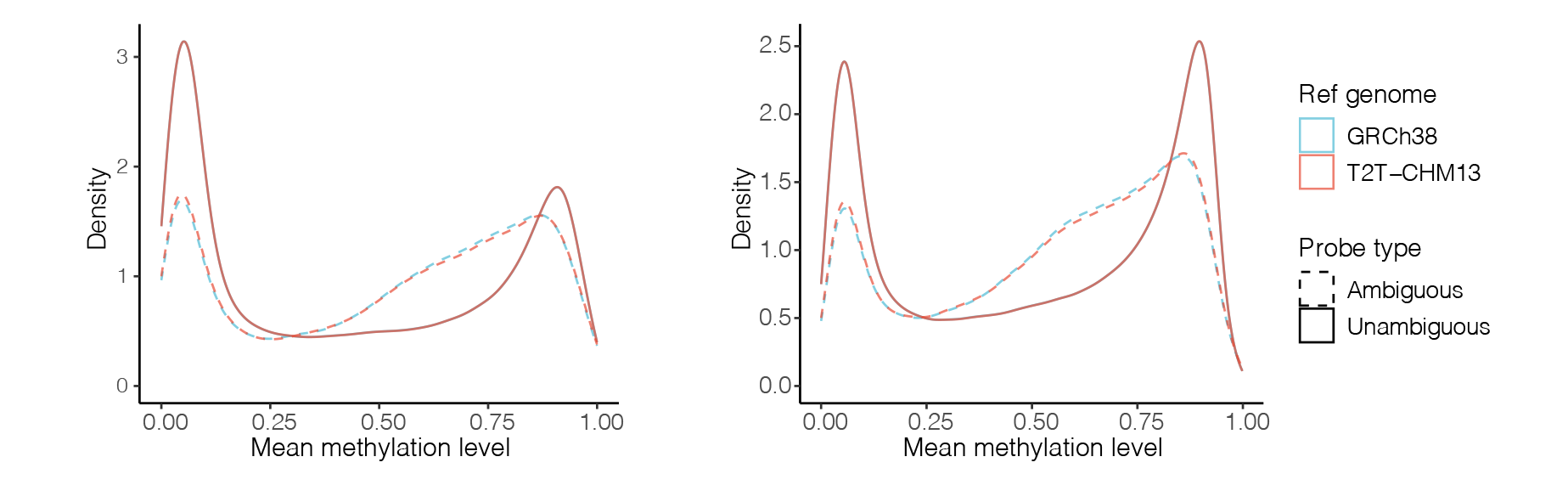

**Figure S7. Mean methylation density differences between ambiguous and unambiguous probes in HM450K (left, *n* = 4) and EPIC (right, *n* = 3) across technical replicates from the IMR-90 cell line.** Ambiguous probes were defined as those with cross-reactivity and/or mismatch.
